# Combinatorial metabolic engineering platform enabling stable overproduction of lycopene from carbon dioxide by cyanobacteria

**DOI:** 10.1101/2020.03.11.983833

**Authors:** George M. Taylor, John T. Heap

## Abstract

Cyanobacteria are simple, efficient, genetically-tractable photosynthetic microorganisms representing ideal biocatalysts for CO_2_ capture and conversion, in principle. In practice, genetic instability and low productivity are key, linked problems in engineered cyanobacteria. We took a massively parallel approach, generating and characterising libraries of synthetic promoters and RBSs for the cyanobacterium *Synechocystis*, and assembling a sparse combinatorial library of millions of metabolic pathway-encoding construct variants. Laboratory evolution suppressed variants causing metabolic burden in *Synechocystis*, leading to expected genetic instability. Surprisingly however, in a single combinatorial round without iterative optimisation, 80% of variants chosen at random overproduced the valuable terpenoid lycopene from atmospheric CO_2_ over many generations, apparently overcoming the trade-off between stability and productivity. This first large-scale parallel metabolic engineering of cyanobacteria provides a new platform for development of genetically stable cyanobacterial biocatalysts for sustainable light-driven production of valuable products directly from CO_2_, avoiding fossil carbon or competition with food production.

## Introduction

Future sustainable economies consistent with net zero greenhouse gas emissions and limiting climate change will require radical changes in global carbon flows, including the development and large-scale deployment of technologies which capture and recycle carbon. Cyanobacteria are the simplest and most genetically-tractable organisms capable of oxygenic photosynthesis, using CO_2_ and sunlight as their sole carbon and energy sources, respectively. Their photosynthetic yield^1^ and growth rate^2^ are similar to fast-growing microalgae, and greater than terrestrial plants^3,4^. Therefore, cyanobacteria have great potential to serve as whole-cell biocatalysts in light-driven, carbon-negative bioprocesses converting atmospheric or waste CO_2_ to products of interest, while avoiding competition with the food chain.

Cyanobacteria have been genetically modified to synthesise a wide range of non-native compounds ranging from commodity chemicals and biofuels to high-value products, including alcohols^5–8^, organic acids^8–11^, diols^12,13^, terpenoids^14–20^, sugars^21–23^ and many others^24–29^. However, production using cyanobacteria is not yet commercially competitive^30–32^ with production by petrochemical or conventional heterotrophic biotechnology approaches, both of which benefit from the current low economic costs of using fossil carbon and emitting CO_2_. There are also issues with cyanobacterial production itself, including lower growth rates and cell densities than many heterotrophs, self-shading^33^ and the crucial genetic and metabolic design challenges of low productivity^34^ and genetic instability^35^. Importantly, these are linked: genetic instability and loss of metabolic pathways causes loss of productivity^8^.

Heterologous DNA often confers a fitness cost on host cells, through the activity of expressed heterologous proteins, or the demands of their synthesis, or both^36^. In the case of heterologous metabolic pathways, the diversion of central metabolites away from endogenous biosynthesis towards non-native products can limit growth. The relative expression of enzymes in heterologous pathways may also be poorly balanced, leading to accumulation of intermediates, which can be toxic^37,38^. Heterologous metabolic pathways which impair growth are genetically unstable, because selection will favour cells in which the pathways are inactivated by spontaneous point mutations or deletions, until such cells predominate. This effect is expected in all organisms, but appears to be quite pronounced in cyanobacteria^35^, perhaps due to the lengthy incubations during strain construction (required to allow genetic segregation, so polyploid strains become homozygous) and cultivation, which provide more opportunity for mutations to occur and to be selected^8,29^. Furthermore, heterologous genes have often been strongly overexpressed in cyanobacteria by inserting them downstream of a strong native promoter^5,9,39–42^, which is unlikely to be optimal and probably contributes to instability^36,43^.

In principle, optimal target concentrations of each enzyme in a pathway could be predicted by metabolic modelling, although this is a substantial undertaking. Furthermore, it is not currently possible to reliably design an individual DNA sequence to produce a set of several enzymes at specific target concentrations, due to the limited predictability of the impact of combinations of different promoters, ribosome-binding sites (RBSs) and codon usage on protein expression, particularly in different sequence contexts^44^ and in lesser-studied organisms. Fortunately, modular DNA assembly provides an alternative solution, by enabling the efficient construction of large libraries of variants of pathway-encoding constructs. By systematically varying the expression of each enzyme combinatorially, it is typically possible to identify pathway variants which perform well by screening. The same assembly methods allow for optimisation by rational replacement of individual parts to fine-tune expression. This combinatorial approach to construction and optimisation of metabolic pathway-encoding constructs is a recent development, and while its effectiveness has been clearly demonstrated in model organisms like *Escherichia coli* and *Saccharomyces cerevisae*^*37,38,45,46*^, it is not yet standard practice, and in many other organisms including cyanobacteria it has not yet been established.

Here we develop a platform for individual and combinatorial assembly and optimisation of metabolic pathway-encoding constructs in cyanobacteria and apply it to light-driven overproduction of lycopene from atmospheric CO_2_ in *Synechocystis* sp. PCC 6803. The platform uses the organism-independent Start-Stop Assembly system, and incorporates synthetic promoters and synthetic RBSs we generated and characterised in *Synechocystis*, along with terminators we validated previously^47^. Lycopene is both a valuable product and a useful model for the challenges of producing terpenoids in photoautotrophs, which is complicated by potentially deleterious competition for common precursors with the biosynthesis of chlorophyll and photoprotective carotenoids. A large combinatorial library of lycopene overproduction pathway variants was constructed and used to study stability and production characteristics. Genetically stable lycopene overproducers were readily obtained. This platform represents a generally-applicable solution to the genetic instability of heterologous metabolic pathways in cyanobacteria, and opens the way to sustainable light-driven production of valuable products directly from CO_2_.

## Results

### A platform for assembly and optimisation of metabolic pathways in cyanobacteria

To develop a general-purpose platform for assembly and optimisation of metabolic pathways in cyanobacteria we required both an efficient multi-part DNA assembly method and sets of expression control parts (promoters, RBSs and transcriptional terminators) with suitable properties. Several DNA assembly methods are suitable for combinatorial assembly^48^, but most are designed for a specific organism. Start-Stop Assembly^38^ offers an ideal general-purpose DNA assembly system, with a unique streamlined assembly hierarchy which typically requires construction of only one new vector to apply the system to a new organism. Therefore we constructed a new Level 2 Start-Stop Assembly vector pGT270 (Figure S1) for the cyanobacterium *Synechocystis* sp. PCC 6803 (hereafter referred to as *Synechocystis*).

At the start of this study there was a lack of promoter and RBS parts suitable for controlling and tuning gene expression in cyanobacteria (although a few have since been published^49,50^). Therefore we set out to generate and screen synthetic libraries in order to identify sets of these expression control parts providing an evenly-distributed, wide range of strengths suitable for controlling and tuning expression levels in cyanobacteria.

A synthetic promoter library (SPL) was designed by the method of Jensen and Hammer^51^, which unlike classic approaches^52–54^ does not mutate the consensus sigma factor-binding -35 or -10 regions, but instead randomises the sequences surrounding these core elements. This is a proven method for generating libraries of synthetic promoters which span an evenly-distributed, wide range of strengths^51,55,56^. A total of 33 bp surrounding the -35 (TTGACA) and -10 (TATAAT) elements of a Type 1 (SigA σ^70^-dependent) *Synechocystis* promoter^57^ (Figure 1b) were randomised using degenerate primers (giving a library size of 7.4×10^19^) and introduced in front of a yellow fluorescent protein CDS (*eyfp*) reporter in the *E. coli*-*Synechocystis* vector pATM2 (Figure 1a). *E. coli* was transformed with the SPL then one hundred transformant colonies were picked (70 randomly and 30 spanning a range of fluorescence levels by visual inspection) and a screening cascade of flow cytometry and sequencing was used for validation (Figure 1a). Promoter clones with unintended mutations or which were non-functional or multimodal (showing sub-populations with differing fluorescence) in *E. coli* were discarded, resulting in a set of 37 promoter clones (Table S1).

**Figure 1.**
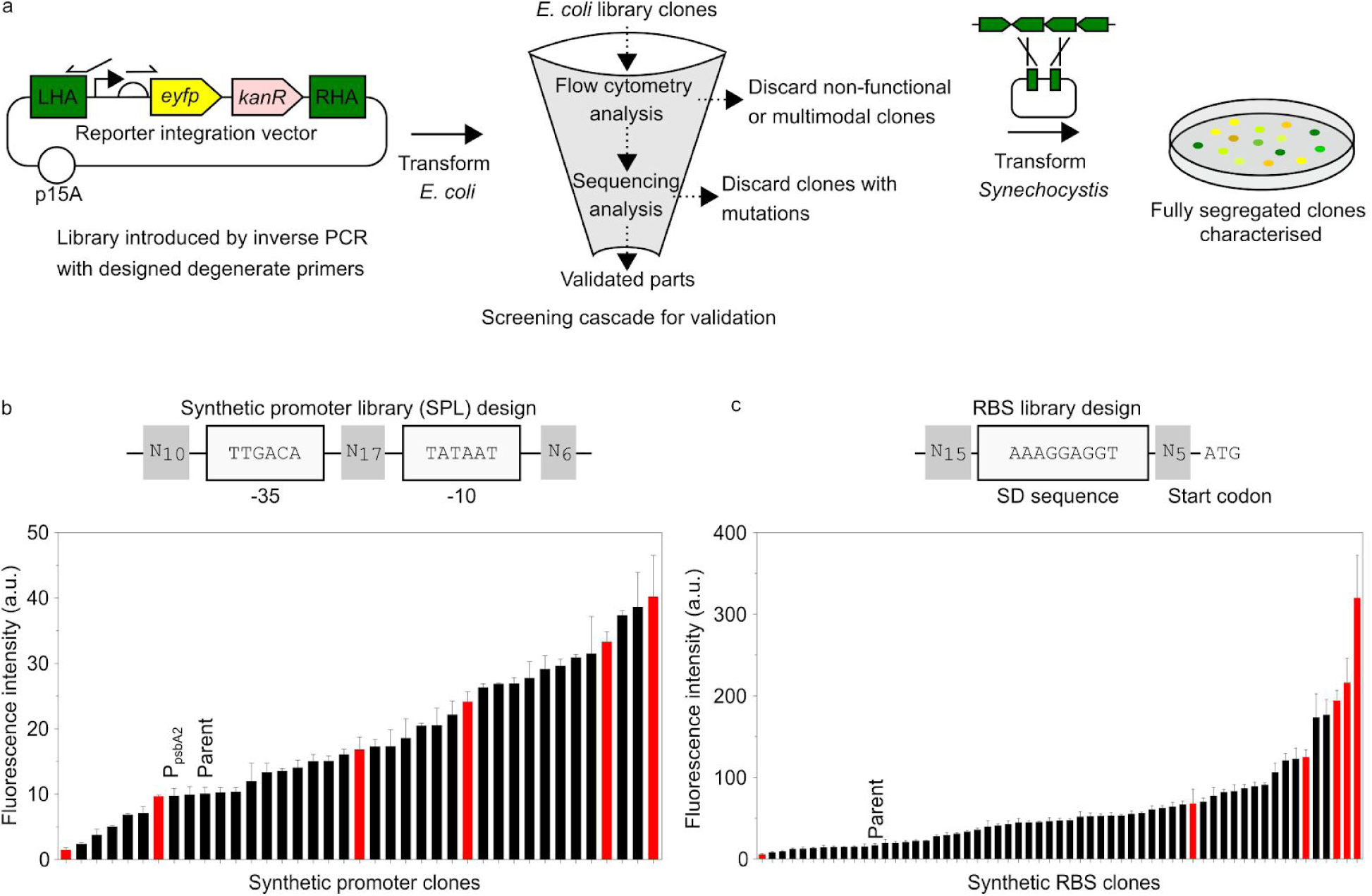
Parts enabling rational and combinatorial pathway assembly in cyanobacteria. (a) Workflow of library design, generation and characterisation. Libraries were generated by inverse PCR and *E. coli* was transformed with each library. Clones were picked (70 randomly and 30 spanning a range of fluorescence levels by visual inspection) and a screening cascade of flow cytometry and sequencing was used for validation. Clones were discarded if their fluorescence was multimodal or weaker than pATM2 (-ve control) or if they contained unintended mutations. *Synechocystis* was transformed and fully segregated before characterisation. (b) Synthetic promoter library (SPL) design conserved -35 and -10 consensus sequences, and randomised the surrounding sequences. *Synechocystis* SPL clones were characterised by flow cytometry (detail in Figure S8). A subset of six promoters was chosen (red bars). ‘Parent’ plasmid pATM2 was used as template to generate SPL. (c) RBS library design conserved the Shine-Dalgarno sequence, and randomised the surrounding sequences. *Synechocystis* RBS library clones were characterised by flow cytometry (detail in Figure S9). A subset of six RBSs was chosen (red bars). ‘Parent’ plasmid pGT77 was used as template to generate RBS library.

*Synechocystis* was transformed separately with each promoter clone plasmid then transformants were subcultured until completely segregated (homozygous). The fluorescence of these 37 segregated *Synechocystis* strains was determined by flow cytometry of mid-linear growth phase cultures under photoautotrophic conditions. We observed a wide distribution of promoter strengths with small increments of strength (Figure 1b), which ranged from very weak (SPL clone 115) to 27-fold stronger (SPL clone 25). Interestingly, over 80% of the synthetic promoter clones were stronger than the positive control strong native cyanobacterial promoter P_psbA2_^58^ (pATM10), and the strongest synthetic promoter SPL clone 25 was three times stronger. The 37 promoter clones were also characterised in *E. coli* where a wide distribution of promoter strengths was again observed (Figure S2), although interestingly no correlation (*R*^2^ = 0.05) was observed between promoter strengths in *Synechocystis* and *E. coli* (Figure S3).

Next we set out to generate RBSs. In principle, computational tools such as the RBS Calculator^59^ can be used to design specific RBS sequences with desired strengths tailored to a specific CDS. In practise, the published RBS design tools remain limited in their predictive ability, particularly for lesser-studied organisms such as cyanobacteria^50,60^. Furthermore, for combinatorial assembly, prediction of the strengths of individual RBSs is less important than the diversity of strengths. Therefore we chose a degenerate approach to design a synthetic RBS library, similar to the promoter library.

In order to obtain RBSs with a wide range of strengths, a series of nine RBS libraries was generated (Figure S4) each differing only in the number of degenerate positions, from 20, giving a library size of 1.1×10^12^ (RBS library 1) to 29, giving a library size of 2.9×10^17^ (RBS library 9). The nine RBS libraries were introduced between a mid-strength promoter (SPL clone 3) from the promoter library and the *efyp* CDS in the *E. coli*-*Synechocystis* vector pGT77 (Figure 1a). *E. coli* was transformed with the nine RBS library assembly products and the fluorescence distributions among populations of cells containing each RBS library were compared in *E. coli* by flow cytometry (Figure S4). A wide range of fluorescence intensities was observed in the RBS libraries with fewest degenerate positions, but as the number of degenerate positions increased, the range of fluorescence intensities narrowed, shifting the population towards that of the negative control (pATM2). 100 transformant colonies were picked (70 randomly and 30 spanning a range of fluorescence levels by visual inspection) from RBS libraries 1, 4 and 9 and initially assessed in singlicate using the same screening cascade as before (Figure 1a) resulting in a set of 58 RBS clones (Table S2), which were then characterised in triplicate, showing a wide distribution of strengths (Figure S5).

*Synechocystis* was transformed separately with each RBS clone and the resulting strains were characterised by flow cytometry as before. We observed a wide distribution of fluorescence intensities that increased in small increments (Figure 1c) from very weak (RBS clone 249) to very strong (RBS clone 4) over a 152-fold range. A very weak correlation (*R*^2^ = 0.34) was observed between RBS strengths in *Synechocystis* and *E. coli* (Figure S6). Interestingly, the 58 RBS clones showed no correlation (*R*^2^ = 0.153) between their measured fluorescence and their predicted ΔG_tot_ in either *Synechocystis* (*R*^2^ = 0.153) or *E. coli* (*R*^2^ = 0.149) (Figure S7), as calculated by the reverse engineering mode of the RBS Calculator v2.0^59^, which is consistent with previous reports^50,60^.

Using the entire set of 37 promoters and 58 RBSs would give up to 2146 expression levels per CDS, which would be excessive and redundant. Instead we chose a subset of six promoters and six RBSs evenly spanning the range of available strengths (Figure 1b and 1c, red bars) with the aim of generating a total of 36 diverse expression levels per CDS, allowing broad sampling of design space.

### Rationally-designed individual pathway constructs were genetically unstable

The promoters, RBSs and Start-Stop Assembly represent a platform for assembly of metabolic pathways in cyanobacteria. To test this platform, and to tackle genetic instability, overproduction of the terpenoid lycopene was adopted as a model system (Figure 2). Lycopene is a valuable product used in the pharmaceutical, food and cosmetic industries^61^ with a strong red colour which can be used as a measure of pathway activity^37,38,62^ including by visual inspection of colony colour, at least in *E. coli*. Lycopene is produced endogenously by *Synechocystis* and other cyanobacteria as a precursor of other carotenoids used for photoprotection, but does not naturally accumulate (Figure 2). To overproduce lycopene a pathway-encoding construct was designed using homologs of three native CDSs (*dxs, crtE*, and *crtB*) to boost pathway flux (*dxs* is a known rate-limiting reaction in both *E. coli*^*37,63,64*^ and cyanobacteria^14,15,65^) and a heterologous CDS (*ctrI*), which provides a bacterial-type one-step desaturase to bypass the native photoautotroph-type two-step desaturase pathway^66^ (*crtP* and *crtQ*) in the conversion of phytoene to lycopene (Figure 2).

**Figure 2.**
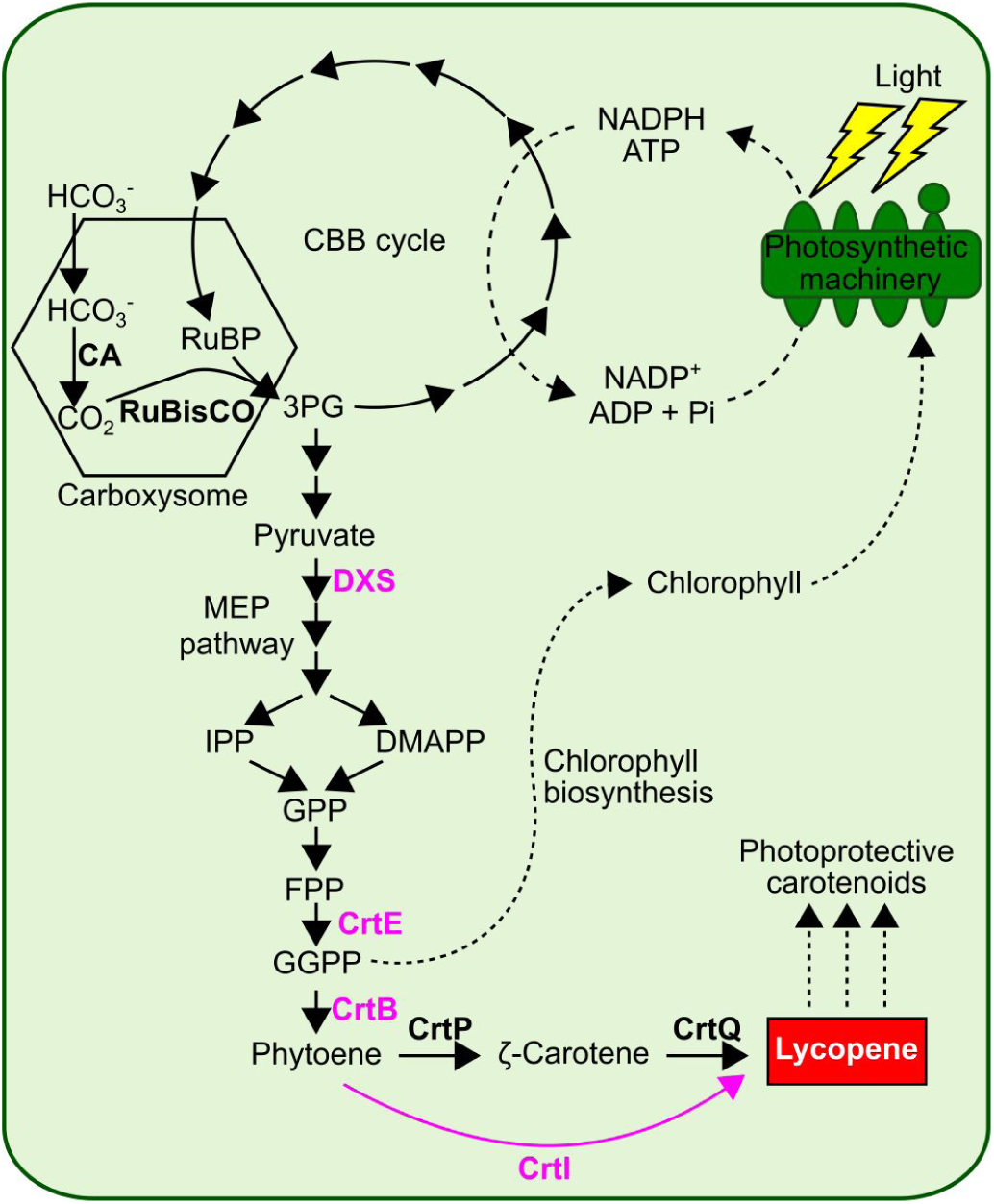
Metabolic design of lycopene overproduction pathway in *Synechocystis*. The lycopene overproduction construct encodes four enzymes (shown in pink). DXS (1-deoxy-D-xylulose 5-phosphate synthase) catalyses the synthesis of 1-deoxy-D-xylulose 5-phosphate from pyruvate and glyceraldehyde 3-phosphate. FPP (Farnesyl diphosphate) is synthesised via the native metabolism of *Synechocystis* (shown in black). GGPP (Geranylgeranyl diphosphate) is synthesised by CrtE (Geranylgeranyl diphosphate synthase) which catalyses the condensation between FPP (Farnesyl diphosphate) and IPP (Isopentenyl diphosphate). CrtB (phytoene synthase) then catalyses the condensation of two GGPP molecules to generate phytoene. Finally, CrtI (phytoene desaturase) catalyses the synthesis of lycopene (shown in red box) from phytoene, bypassing the native two-step desaturase pathway (CrtP and CrtQ) via ζ-carotene.

We designed four individual lycopene pathway-encoding constructs for *Synechocystis* using different design strategies, to test their effects on genetic stability, aiming to identify a strategy allowing stable expression and continued overproduction. The first design (pGT432) used a traditional strong overexpression rationale with the strongest promoter and RBS driving expression of each CDS. The second design (pGT433) aimed to avoid excessive expression by using medium-strength promoters and RBSs for all four CDSs. The third design (pGT434) followed recently-described principles for minimising metabolic burden, which have been demonstrated in *E. coli*^*36*^. This design uses a strong promoter and weak RBS to reduce ‘overinitiation’ of translation, because high translation is more stressful for the cell than high transcription. As a high-burden control, the fourth pathway design (pGT435) reversed this principle, using a weak promoter and strong RBS.

We opted to express each CDS in a monocistronic configuration (with its own promoter and terminator) to maximise their independence, and therefore included four terminators from a set we recently characterised in *Synechocystis*^*47*^. The parts (promoters, RBSs, CDSs and terminators) were each stored and verified in Start-Stop Assembly format and the four pathways (pGT432-435) were individually assembled by hierarchical Start-Stop Assembly (Figure 3 and Figure S10). After three independent attempts it was not possible to assemble the supposedly low-burden design pGT434, suggesting that unexpectedly it is toxic to *E. coli*. The other three designs were successfully assembled, giving red-coloured *E. coli* colonies showing lycopene production.

**Figure 3.**
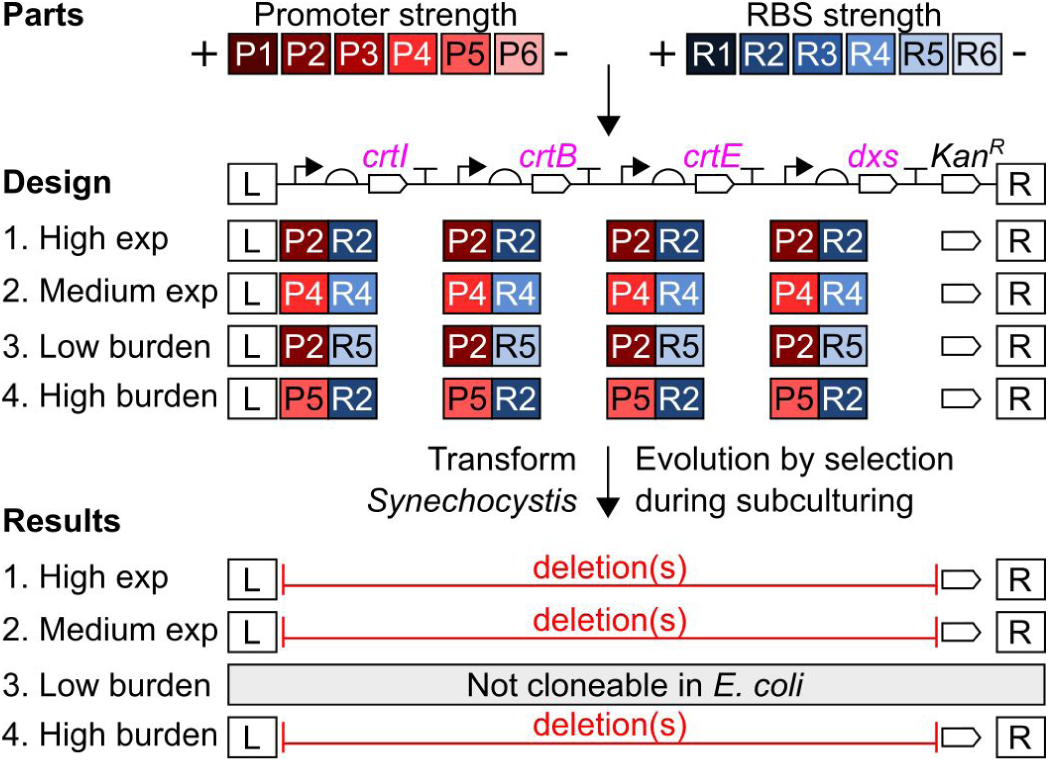
Rational design and genetic instability of individual lycopene overproduction constructs for *Synechocystis*. Four individual pathway constructs were designed: high expression (pGT432), medium expression (pGT433), low burden (pGT434) and high burden (pGT435). Each construct was individually assembled using Start-Stop Assembly in the destination vector pGT270 (Figure S10). pGT434 was not cloneable in *E. coli* (three independent attempts). *Synechocystis* was individually transformed with each pathway and PCR screening was used to assess genetic stability (detail in Figure S11). In each case flanking sequences were present and antibiotic selection pressure maintained the kanamycin resistance gene *kan*^*R*^, but gross deletions in the lycopene pathway-encoding constructs were observed. Promoters in descending order of strength as characterised using EYFP: P1 = SPLc25, P2 = SPLc19, P3 = SPLc47, P4 = SPLc3, P5 = SPLc45 and P6 = SPLc17. RBSs in descending order of strength as characterised using EYFP: R1 = RBSc4, R2 = RBSc21, R3 = RBSc110, R4 = RBSc123, R5 = RBSc48 and R6 = RBSc249.

To assess the high expression (pGT432), medium expression (pGT433) and high burden (pGT435) lycopene pathway constructs, *Synechocystis* was transformed separately with each plasmid and serially subcultured under antibiotic selection in an attempt to obtain homozygous clones. Individual transformant colonies grew more slowly than usual, appearing after approximately two weeks instead of one, suggesting a burdensome effect of the pathways. After four subcultures we assessed genetic stability (Figure 3 and Figure S11) and found that all three remaining lycopene pathway designs had acquired gross deletions which had become homozygous, presumably due to selection pressure acting against the burden imposed by the constructs. Therefore, due to genetic instability either in *E. coli* or *Synechocystis*, none of the four individual pathway designs was suitable for sustained overproduction of lycopene in *Synechocystis*, so an alternative approach was needed.

### Combinatorial design overcomes trade-off between productivity and genetic instability

Next we explored a combinatorial approach to address the genetic instability observed with the individually-designed pathways. Using the same assembly system, monocistronic architecture and gene order as before, we designed a library of lycopene pathway encoding constructs in which the promoter and RBS of each CDS was varied combinatorially. The six promoters and six RBSs identified earlier were preassembled as composite promoter-RBS parts, of which 34 of 36 possible combinations were cloneable, giving a maximum combinatorial library size of 1.3×10^6^. The pathway library (pGT547) was generated using hierarchical Start-Stop Assembly with suitable mixtures of parts used in place of individual parts in order to introduce the combinatorial diversity at Level 1, and to propagate the diversity to the complete pathways at Level 2 (Figure 4, Figure S12). From the *E. coli* transformation, 20 clones (pGT437-456) were picked at random and 20 clones (pGT457-476) were chosen by prescreening 96 (randomly-picked) clones for their lycopene production in *E. coli*. Of the 20 chosen by prescreening, ten were the strongest lycopene producers and the other ten were evenly-distributed throughout the observed range of lycopene concentrations (Figure 4 and Figure S13).

**Figure 4.**
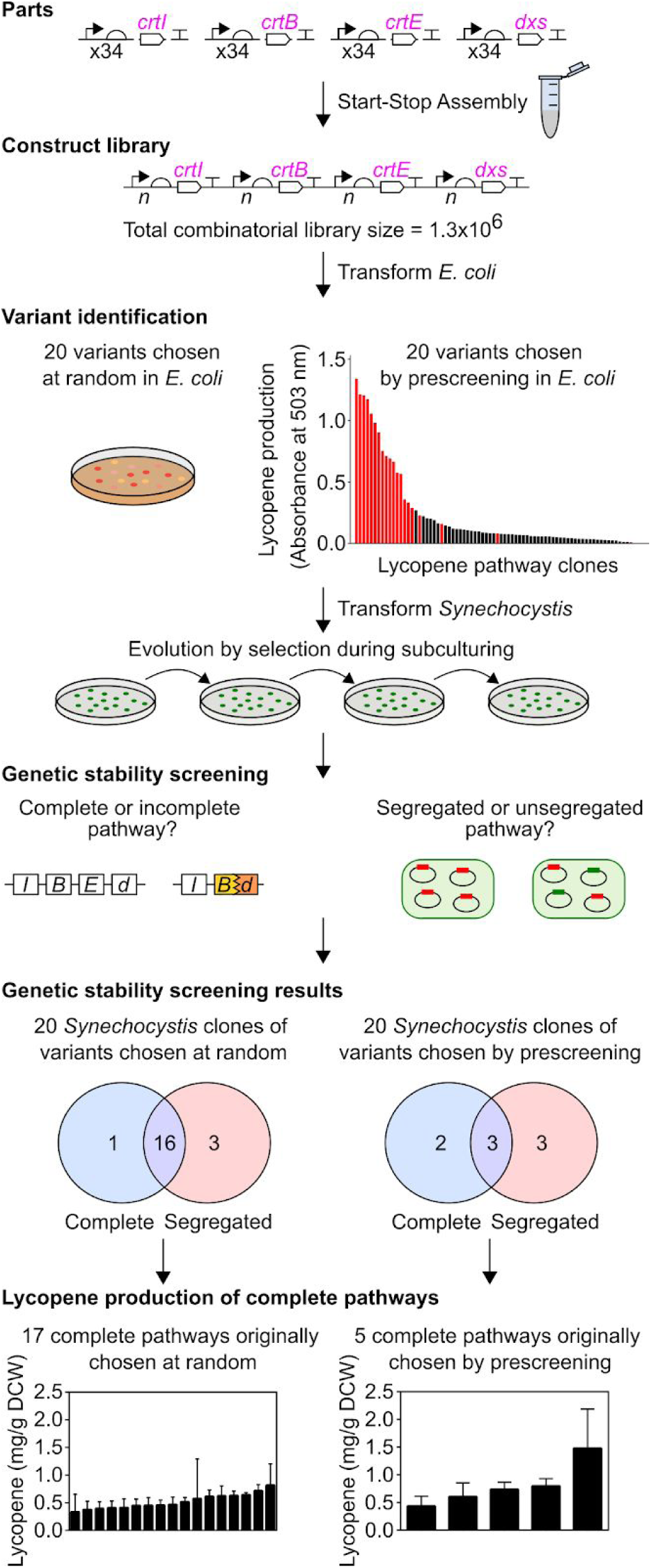
Genetic design, selection and evaluation of combinatorial lycopene pathway library. A library of lycopene pathway construct variants was assembled using Start-Stop Assembly in the destination vector pGT270. For combinatorial assembly equimolar mixtures of 34 composite promoter-RBS parts labelled ‘x34’ were used in Level 1 assemblies (Figure S12). The uncertain representation of each promoter and RBS at each position following assembly is represented by ‘*n*’. The maximum library size was 34^4^ = 1.3×10^6^. *E. coli* was transformed with the combinatorial library and pathway clones were either chosen at random or after prescreening pathway clones performance in *E. coli* (see text). *Synechocystis* was transformed with the chosen pathway variants, after which PCR screening was used to assess the completeness and segregation of the resultant clones (PCR screening details shown in Figure S11). *Synechocystis* clones containing complete pathway constructs were grown in photoautotrophic batch cultures with constant light for two weeks, then lycopene production was measured (detail in Figure S15). Lycopene concentration was determined as mg lycopene per g dry cell weight (DCW). The error bars shown represent the standard deviation of three independent biological replicates.

*Synechocystis* was transformed separately with each of the 40 library variant plasmids and subcultured under antibiotic selection. As previously observed, transformant colonies grew more slowly than usual, suggesting metabolic burden, and after four subcultures we assessed genetic stability in terms of completeness and segregation as before (Figure 4). Interestingly, 16 of the 20 randomly-chosen pathway variants were genetically stable (complete and segregated) in *Synechocystis* (Figure 4), whereas only three of the 20 pathway variants chosen by prescreening in *E. coli* were stable (Figure 4), as the other 17 contained deletions and/or failed to segregate.

The lycopene production of the 22 *Synechocystis* clones containing complete pathway constructs was determined using small-scale batch cultures grown photoautotrophically for two weeks (Figure 4). Lycopene accumulation was detected in all 22 clones (unlike the wild type) but only small differences in lycopene production were observed among the pathway variants, ranging between 0.3-1.5 mg/g DCW (Figure 4). Interestingly, the same 22 pathway variants gave considerably greater diversity of lycopene production in *E. coli*, ranging between 0-40 mg/g DCW (Figure S14).

### Promoter and RBS preferences are associated with coding sequences, not genetic stability

To assess the impact of the combinatorial design, the 40 pathway variants used to transform *Synechocystis* were sequenced. All expected parts were present in multiple variants. Some deletions or misassemblies could be identified (Figure S17 and S18). As diversity of expression was introduced using the same mixture (the exact same tube) of promoters and RBSs for each CDS during the assembly (Figure S12), any substantial differences in frequencies of parts among the complete pathway constructs may reflect biological selection rather than assembly bias (Figure 5 and Figure S19). There were no obvious differences in part frequencies between the segregated and non-segregated variants, one of the key aspects of genetic stability. In contrast, part frequencies clearly differ among the four CDSs, for both segregated and non-segregated variants. For example, we observed all six RBSs for *crtB* whereas for *dxs* we only observed three of the six RBSs. This observation is consistent with biological selection, which has presumably led to a preference towards certain expression levels.

**Figure 5.**
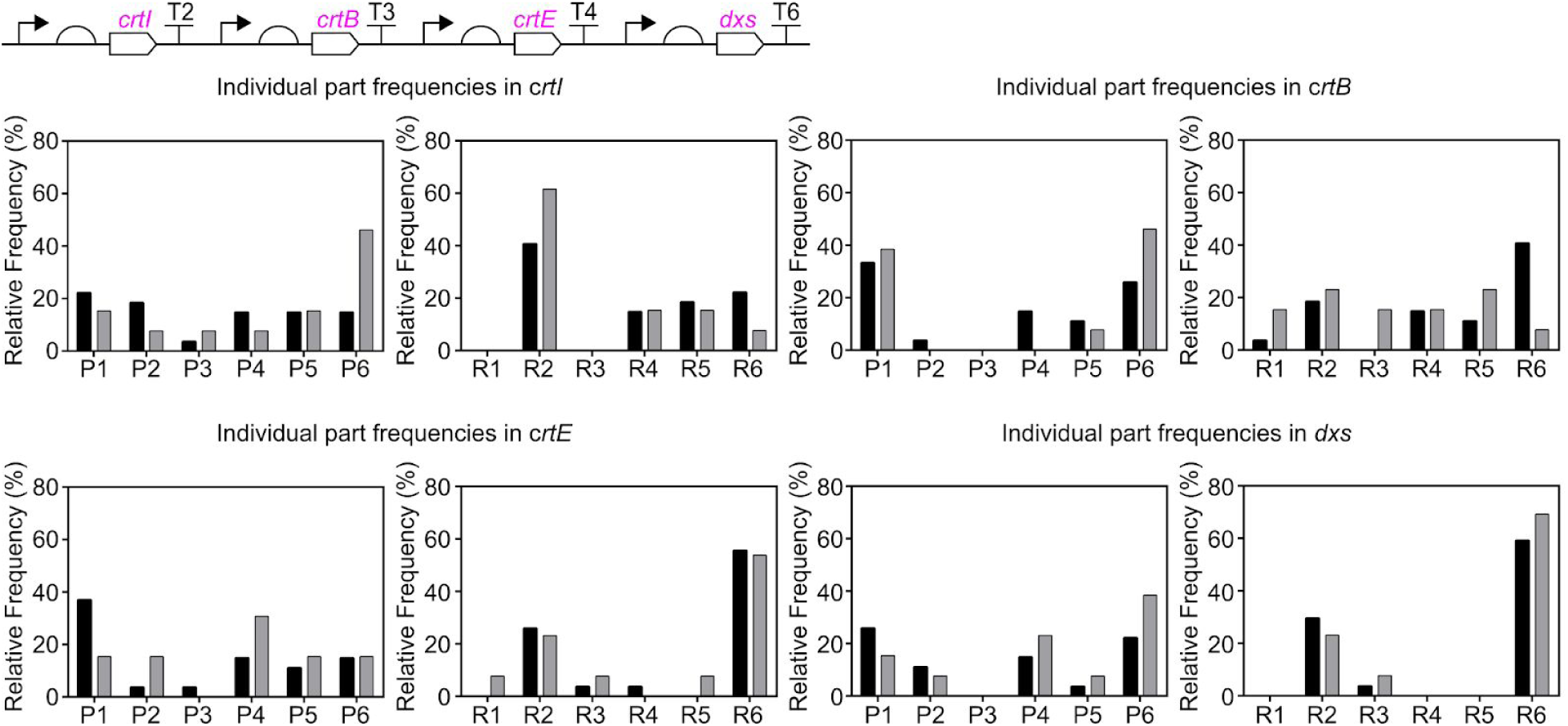
Frequencies of promoter and RBS parts in pathway variants. All 40 pathway variants used to transform *Synechocystis* were sequenced and the promoter and RBS upstream of each CDS were identified. Relative frequencies of promoter and RBS parts: In segregated pathway variants, black bars; in non-segregated pathway variants, grey bars. Promoters, in descending order of strength as characterised using EYFP: P1 = SPLc25, P2 = SPLc19, P3 = SPLc47, P4 = SPLc3, P5 = SPLc45 and P6 = SPLc17. RBSs, in descending order of strength as characterised using EYFP: R1 = RBSc4, R2 = RBSc21, R3 = RBSc110, R4 = RBSc123, R5 = RBSc48 and R6 = RBSc249.

### Increasing precursor supply enhances lycopene production

Glucose supplementation has been used to increase the supply of precursors to product-forming pathways^67^, so we tried this approach to increase lycopene overproduction by the engineered strains, and to test the hypothesis that the low diversity of lycopene overproduction in *Synechocystis* might be caused by limited precursor availability. We compared the lycopene production of low (pGT439), medium (pGT455) and high (pGT462) producing strains (Figure S16) in photoautotrophic (light, no glucose) and mixotrophic (light and glucose) conditions which boost precursor supply^67^ (Figure 6). Supplementary glucose successfully increased overproduction by the medium and high producing strains (Figure 6). The modest level of increase suggests that the low diversity of lycopene overproduction in the library is not caused by limited precursor availability, and may instead be due to increased flux from lycopene to other terpenoids or low tolerance to changes in production of these physiologically important products.

**Figure 6.**
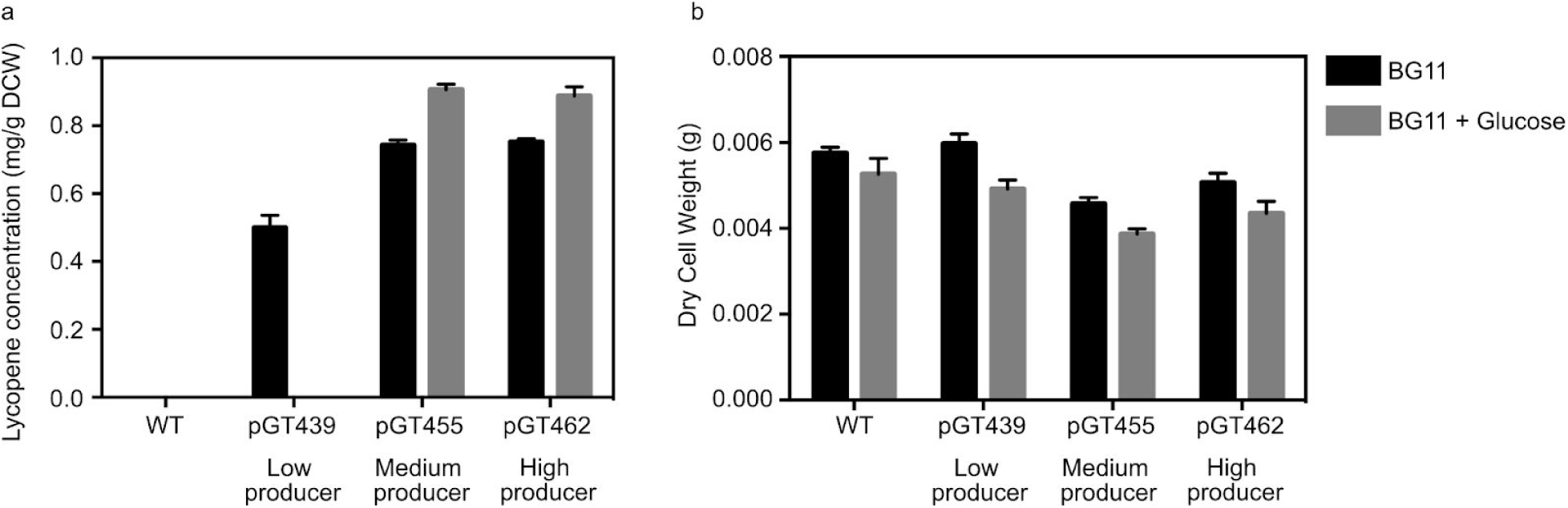
Effect of glucose supplementation on lycopene overproduction. The low, medium and high producing *Synechocystis* clones were grown in photoautotrophic conditions (light, no glucose; black bars) or mixotrophic conditions (light and glucose; grey bars) in batch cultures for two weeks with constant light, then (a) lycopene production and (b) dry cell weight (DCW) were determined. Error bars represent the standard deviation of three independent biological replicates.

## Discussion

Genetic stability is essential for industrial microorganisms, so the reported genetic instability of genetically modified cyanobacteria^35^ is a key challenge for cyanobacterial synthetic biology^8,29,68–71^. Despite its importance, genetic instability of heterologous DNA in cyanobacteria has not been systematically studied. Several aspects of the present study provide new insight into this problem. Firstly, we used a large-scale combinatorial approach to metabolic pathway construction. Such approaches have proven useful for simultaneously optimising expression of multiple genes in other organisms^37,38,45,46^, but have not been established in cyanobacteria. Only two small-scale examples of combinatorial designs have been reported, the first varying a small number of alternative enzyme-encoding sequences^60^ and the second varying only the RBSs in a two-enzyme pathway^72^. In both cases, the total number of variants constructed was very small (14-25 variants) compared to the libraries of millions of variants described in *E. coli* and *Saccharomyces cerevisae*^37,38,46^. Here, construction of a large library (of over a million variants) and evaluation of numerous variants (rather than the typical one or two) provided more scope to overcome the low tolerance to heterologous genes and/or poor control of expression presumably involved in reported examples of genetic instability. Secondly, overproduction of the target product lycopene is likely to be sensitive to genetic instability, and hence able to reveal design issues. Lycopene is a terpenoid, one of the largest and most diverse classes of natural products, with existing or emerging applications in the pharmaceutical, flavour, fragrance and fuel industries^61^. However, modifying production of terpenoids in photoautotrophs is challenging, because the monoterpenoid GGPP is a precursor of carotenoids, chlorophyll, ubiquinones, plastoquinone and phylloquinone^73^. These products are all essential components of the photosynthetic machinery, involved in their function and/or protection^73^, and are therefore essential in photoautotrophs. As a result, the overproduction of any terpenoid competes with the biosynthesis of these essential products. Thirdly, our constructs allowed direct comparisons between *E. coli* and *Synechocystis* to be made at each stage, from the performance of individual promoter and RBS parts, to the relative lycopene overproduction of assembled pathway variants, to their associated genetic stability.

Genetic instability was prevalent in the combinatorial lycopene pathway library, as anticipated. However, many variants were genetically stable, over many generations during subculturing in the transformation process, and yet crucially many of these stable variants were also able to overproduce lycopene. Due to the slower than usual growth observed, this subculturing took approximately eight weeks, which effectively represents a long laboratory evolution experiment, and a substantial window of opportunity for cells to escape burden imposed by variants through spontaneous mutation. Genetic stability despite this long evolution window indicates real robustness and bodes well for the prospect of overproducing other terpenoids.

The synthetic promoters characterised using the fluorescent reporter behaved very differently in *E. coli* and *Synechocystis* (Figure S3). The RNA polymerase of these organisms differs somewhat^74^ although the complete lack of correlation was surprising. The synthetic RBSs did show a modest correlation between *E. coli* and *Synechocystis* (Figure S6), although perhaps weaker than might have been expected given the near-identical anti-Shine Dalgarno sequences of the two organisms^75^. The strengths of these synthetic RBSs were not computationally predictable for either organism (Figure S7). It was therefore to be expected that the pathway variants combinatorially assembled using these parts produced different relative amounts of lycopene in *E. coli* and *Synechocystis* (Figure S20), particularly given the profoundly different physiology, metabolism and growth rates of these organisms^76^. Interestingly, most of the pathway variants which produced large amounts of lycopene during prescreening in *E. coli* were not genetically stable in *Synechocystis* (Figure 4). These apparently conflicting observations might be explained if there is an underlying correlation between the performance of pathway variants in the two organisms, but *Synechocystis* is more sensitive to metabolic burden caused by high-expressing or high-producing variants, leading to mutations which in turn prevent higher production being observed. This would be consistent with both the reported genetic instability of recombinant cyanobacteria, and with the expected sensitivity of cyanobacteria to perturbations in the production of lycopene (Figure 3).

This work directly shows that the previously anecdotally-reported genetic instability of heterologous DNA in cyanobacteria is real, but the large fraction of stable overproducers in our sample (80% of variants chosen at random, 15% of those chosen by prescreening in *E. coli*) reveals that the problem can be overcome through suitable genetic design. The lack of association between the frequencies of promoter and RBS parts and the genetic stability of variants containing them (Figure 5) does not provide principles for the rational design of individual genetically stable constructs. Faced with limited predictive ability in a design-build-test-learn (DBTL) cycle framework, improvements could be obtained either by accelerating the cycles or taking highly parallel approaches^46^. CRISPR-Cas9 has been used to accelerate segregation of recombinant cyanobacteria^77^, and chromosomal integration can be replaced by replicative plasmids, although these are inherently unstable^8^. In this first cyanobacterial example of a highly parallel approach to strain engineering, pathway construct variants with the desired characteristics (genetic stability combined with overproduction) were obtained in a single round. This is particularly noteworthy and useful in relatively slow-growing organisms like cyanobacteria, in which multiple iterative DBTL cycles could be prohibitively time-consuming.

Although previous studies have not directly dissected genetic instability in recombinant cyanobacteria, the problem is indirectly addressed by an emerging body of research on coupling production of target compounds with growth, in order to generate a selection pressure for sustained production^78,79^. Such selection can only act on the genetic variants it encounters, so if used with suboptimal conventional individual pathway designs, the growth coupling strategy could force cells to maintain burdensome pathway-encoding constructs, leading to poor performance. However, together with the combinatorial approach reported here, efficient systems might be developed, giving optimal performance in terms of productivities and yields, along with long-term stability.

This study reveals simple principles for rational design of combinatorial libraries of pathway-encoding constructs, from which individual variants with desired properties can be obtained by screening a modest number of variants: Synthetic promoter and RBS parts are generated, characterised and used to implement a broad, sparse sampling of design space with few prior assumptions. The parts and vectors needed for this platform are available to the community from Addgene, and should allow cyanobacterial synthetic biology to much more effectively tackle the genetic instability “elephant in the room”^35^ and develop stable cyanobacterial biocatalysts for sustainable light-driven production of valuable products directly from CO_2_.

## Methods

### Bacterial strains and growth conditions

*E. coli* strain DH10B was used for all DNA assembly, all other cloning, and all other experiments described in this study. *E. coli* DH10B cells were routinely grown in LB medium (tryptone 10 g l^−1^, yeast extract 5 g l^−1^ and NaCl 5 g l^−1^) at 37 °C, with shaking at 225 rpm, or on LB agar plates (containing 15 g l^−1^ bacteriological agar) at 37 °C. LB was supplemented with ampicillin (100 µg ml^−1^), tetracycline (10 µg ml^−1^) or kanamycin (50 µg ml^−1^) as appropriate. *E. coli* cells were routinely transformed by electroporation^80^.

*Synechocystis* sp. PCC 6803 (the glucose-tolerant derivative of the wild type, obtained from the Nixon lab at Imperial College London) was routinely grown in TES-buffered (pH 8.2) BG11 medium (photoautotrophic growth)^81^. Cultures were supplemented with 5 mM glucose as appropriate (mixotrophic growth). Cultures were grown at 30 °C with agitation at 150 r.p.m and in constant white light at 50 μmol m^−2^ s^−1^. BG11 was supplemented with kanamycin (30 μg ml^−1^) as appropriate.

### Construction of synthetic promoter library and synthetic RBS library

Tables of oligonucleotides (Table S3), plasmids (Table S4) and synthetic DNA (Table S5) are provided in the Supplementary Information. The synthetic promoter library and RBS libraries were generated by inverse PCR with 5′-phosphorylated degenerate primers using pATM2 and pGT77 (SPL clone 3) as templates, respectively. After PCR, DpnI was added to the PCR product to digest the template. The PCR product was circularised by ligation with T4 DNA ligase. The ligation product was used to transform *E. coli*. The SPL was generated using the degenerate primers oligoATM3 and oligoATM5. The nine RBS libraries were generated using the degenerate primers oligoATM51-60.

### Construction of *Synechocystis* Start-Stop Assembly destination vector

To generate pGT270, the pSHUTTLE2 backbone was PCR-amplified (oligoGT373 and oligoGT374) and assembled with the Level 2 Start Stop Assembly cassette PCR-amplified from pStA212 (oligoGT375 and oligoGT376) by SOE-PCR. The PCR product was treated with DpnI to digest the template, then circularised by ligation with T4 DNA ligase. The ligation product was used to transform *E. coli*. The plasmids of transformant colonies were purified and sequence verified.

### Storing genetic parts in Start-Stop Assembly storage vector

Composite promoter-RBS parts were cloned in pStA0 to generate pGT287-290, 293-297, 300-303, 305-309, 311-315, 317-321 and 425-430 (Table S6). This was achieved by inverse PCR with 5′-phosphorylated primers (oligoGT436-446 and 583) using pStA0 as a template. After PCR, DpnI was added to the PCR product to digest the template. The PCR product was circularised by ligation with T4 DNA ligase. The ligation products were used to transform *E. coli*. The plasmids of transformant colonies were purified and sequence verified. To store in Start-Stop Assembly format one additional transcriptional terminator validated in *Synechocystis*, pGT424 (pStA0::ECK120015170) was generated by inverse PCR with 5′-phosphorylated degenerated primers (oligoGT579 and oligoGT578) using pStA0 as the template. After PCR, DpnI was added to the PCR product to digest the template. The PCR product was circularised by ligation with T4 DNA ligase. The ligation product was used to transform *E. coli*. The plasmids of transformant colonies were purified and sequence verified.

### Assembly of lycopene pathway constructs

The four individual lycopene pathway constructs (pGT432-435) and the combinatorial lycopene pathway construct library (pGT547) were assembled using Start-Stop Assembly^38^. For each of the four individual constructs and the library, four Level 1 expression units encoding *crtI, crtE, crtB* and *dxs* were assembled from parts or mixtures of parts. These four expression units were then assembled to generate the pathway-encoding constructs in the Level 2 destination vector pGT270. For pGT432, each of the four CDSs were assembled using promoter SPL clone 19 and RBS clone 21 (Figure S10). For pGT433, each of the four CDSs were assembled using promoter SPL clone 3 and RBS clone 123 (Figure S10). For pGT434, each of the four CDSs were assembled using promoter SPL clone 19 and RBS clone 48 (Figure S10). For pGT435, each of the four CDSs were assembled using promoter SPL clone 45 and RBS clone 21 (Figure S10). For the combinatorial library pGT547, an equimolar mixture of the 34 promoter-RBS composite parts (pGT287-290, 293-297, 300-303, 305-309, 311-315, 317-321 and 425-430) was used in Level 1 assemblies of the four CDSs. All white colonies from the transformation plates of the four Level 1 reactions were picked, resuspended as a mixture in P1 buffer and DNA was isolated using a miniprep procedure. The mixtures of Level 1 constructs were then used in the Level 2 assembly reaction to generate the lycopene pathway library (Figure S12).

### *Synechocystis* strain construction

*Synechocystis* cells were grown in BG11 medium and incubated under phototrophic conditions, 30 °C, 150 r.p.m, 50 μmol photons m^-2^ s^-1^ light and incubated to an OD_750_ of 0.5. Following incubation, cells were harvested from 4 ml of culture by centrifugation at 3200 g for 15 min. Pellets were resuspended in 100 μl BG11, then 100 ng of plasmid DNA was added and the mixture was incubated at 100 μmol photons m^−2^ s^−1^ light for 60 min. Cells were transferred onto BG11 plates and incubated at 50 μmol photons m^−2^ s^−1^ light for 24 h at 30 °C. Cells were collected and transferred onto BG11 plates supplemented with 30 μg ml^−1^ kanamycin. To achieve complete segregation, single colonies were subcultured four times on selective BG11 plates, after which segregation was assessed by PCR using primers oligoAH48 and oligoAH49. Completeness of lycopene pathway constructs was assessed using three sets of primers: oligoAH34 and oligoGT603, oligoGT654 and oligoGT605, and oligoGT653 and oligoAH49.

### Flow cytometry analysis of *Synechocystis*

*Synechocystis* cells were grown in 5 ml BG11 medium, supplemented with kanamycin (30 μg ml^−1^), from isolated transformant clones of *Synechocystis* and incubated in 25 ml vented culture flasks under phototrophic conditions, 30 °C, 150 r.p.m, 50 μmol photons m^-2^ s^-1^ light and incubated for 48 h. Following incubation, cultures were diluted to an OD_750_ of 0.1 and incubated for a further 24 hours. Cultures were diluted 1:50 in filtered PBS and were immediately subjected to flow cytometry analysis. A two-laser FACScan flow cytometer (Becton-Dickinson) equipped with an automated multiwell sampler (AMS) was used. The sample was analysed at a data rate of 1000-1500 events s^-1^ for 45 sec. A 488 nm argon ion laser was used for excitation of the EYFP reporter. Detection used a 530 nm band-pass filter (FL1) with a gain of 700 V. All part characterisation assays in *Synechocystis* were conducted using three biological replicates.

### Flow cytometry analysis of *E. coli*

*E. coli* DH10B cells were transformed with purified plasmids. Individual transformant colonies were used to inoculate 200 μl LB broth supplemented with the appropriate antibiotic, in a 96 U-shaped 1.2 ml well plate covered with sterile breathable sealing film (Breathe Easy) and grown at 37 °C at 700 r.p.m for 16 h in a Multitron shaker (Infors-HT). Cultures were subcultured by 1:1000 dilution into 200 μl ml fresh LB broth with the appropriate antibiotic and grown at 37 °C at 700 r.p.m for 4 h. Cultures were diluted 1:50 in filtered PBS and were immediately subjected to flow cytometer analysis. A two-laser FACScan flow cytometer (Becton-Dickinson) equipped with an automated multiwell sampler (AMS) was used. The sample was analysed at a data rate of 1000-2000 events sec^-1^ for 30 sec. A 488 nm argon ion laser was used for excitation of the EYFP reporter. Detection used a 530 nm band-pass filter (FL1) with a gain of 900 V. All part characterisation assays in *E. coli* were conducted using five biological replicates.

### Spectrophotometric quantification of lycopene

*E. coli* DH10B cells were transformed with 5 μl of the combinatorial lycopene pathway library reaction product. Individual transformant colonies were used to inoculate 500 μl LB broth supplemented with the appropriate antibiotics, in a 96 U-shaped 1.2 ml well plate covered with sterile breathable sealing film (Breathe Easy) and grown at 30 °C at 700 r.p.m for 48 h in a Multitron shaker (Infors-HT). Following incubation, the liquid cultures were centrifuged at 4000 r.p.m for 15 minutes before removing the supernatant and adding 100 μl of methanol and 200 μl of hexane to the bacterial cell pellets. After vortexing for 2 minutes, 100 μl of deionised water was added before vortexing the mixture again for 2 minutes. The solution underwent centrifugation at 4000 r.p.m for 20 minutes and the top hexane layer was transferred to a shallow 96-well plate and absorbance at 530 nm was determined (BMG LABTECH POLARstar Omega).

### Growth of *Synechocystis* strains for lycopene production

*Synechocystis* cells were grown in 5 ml BG11 medium, supplemented with kanamycin (30 μg ml^−1^), from isolated transformant clones of *Synechocystis* and incubated in 25 ml vented culture flasks under phototrophic conditions, 30 °C, 150 r.p.m, 50 μmol photons m^-2^ s^-1^ light and incubated for 48 h. Following incubation, cultures were diluted to an OD_750_ of 0.1 and incubated for a further 2 weeks

### Growth of *E. coli* strains for lycopene production

*E. coli* DH10B cells were transformed with purified plasmids. Individual transformant colonies were used to inoculate 5 ml LB media supplemented with the appropriate antibiotic and grown at 30 °C at 225 r.p.m for 24 h. Cultures were subcultured by 1:1000 dilution into 5 ml fresh LB and grown for 48 hours.

### Dry cell weight measurements

Dry cell weight (DCW) was determined by centrifugation of 2 ml of liquid culture at 16,900 g for 5 minutes to remove supernatant, the bacterial cell pellet was washed in deionised water and then centrifuged again and dried at 70 °C until a constant weight was obtained.

### Lycopene extraction

To extract lycopene, 2 ml of liquid culture was centrifuged at 14,000 r.p.m for 5 minutes to remove supernatant. The bacterial cell pellet was washed in deionised water and then centrifuged as before. The bacterial cell pellet was resuspended in 1 ml acetone and incubated at 55 °C for 15 minutes to extract lycopene. The supernatant was obtained by filtration through a 0.22 μm pore-size nylon membrane for LC-DAD analysis.

### LCDAD quantification of lycopene

Lycopene was detected and measured using an Agilent LC system with UV/Vis diode array detector. Absorbance at 450 nm and 471 nm were monitored and the peak area corresponding to each component integrated to provide a measure of abundance. The LC column used was an Acquity UPLC Peptide BEH C18 column (2.1 × 100mm, 1.7 μm, 300Å, Waters). LC buffers were 50% of methanol in water (A) and 25% of ethyl acetate in acetonitrile (B). All the solvents used were HPLC grade. The LC method was 6.5 minutes in total with 1.5 minutes of post run time (0-1 min: 30% A, 70% B; 1-6 min: 0.1% A, 99.9% B; 6-6.5 min: 30% A, 70% B; at a flow rate of 0.3 ml/min). The injection volume for the samples was 1.0 μl. Commercially available lycopene (Sigma-Aldrich) was dissolved in acetone as a standard and a standard curve was generated.

## Acknowledgements

The authors thank Dr Soo Mei Chee for assistance with LC-UV analysis, Antonio Torres-Méndez for assistance with preliminary experimental design, and Professor Tom Ellis, Dr Karen Polizzi and Dr Patrik Jones for critical input. The authors acknowledge funding from the Imperial College London Schrödinger Scholarship Scheme, BBSRC grants BB/M002454/1 and BB/M011321/1, and Imperial College London’s Excellence Fund for Frontier Research.

## Author Contributions

G.T. and J.H. designed the study; G.T. performed experiments; G.T. and J.H. prepared the manuscript.

## Competing Interests statement

None declared.

