## Supplementary Information for "Combinatorial metabolic engineering platform enabling stable overproduction of lycopene from carbon dioxide by cyanobacteria"

**George M. Taylor[1] and John T. Heap*[1,2]**
[1] Imperial College Centre for Synthetic Biology, Department of Life Sciences, Imperial College London, London, SW7 2AZ, United Kingdom
[2] School of Life Sciences, The University of Nottingham, University Park, Nottingham, NG7 2RD, United Kingdom

*** Corresponding author**
John T. Heap,

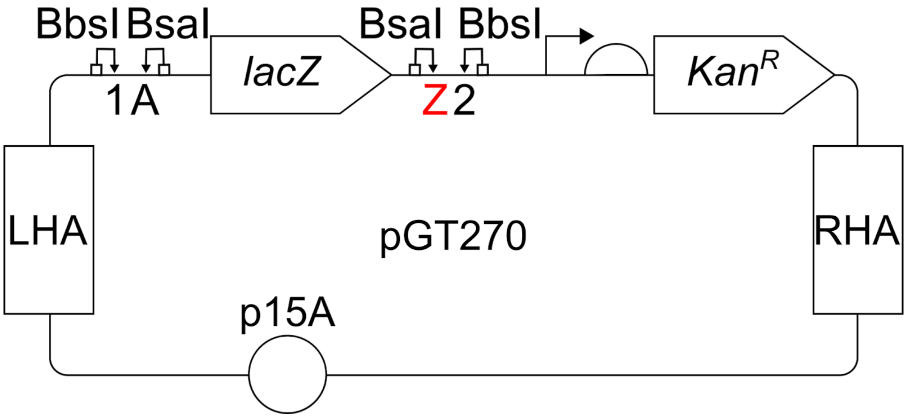


**Figure S1. *E. coli-Synechocystis* shuttle vector pGT270 containing Level 2 Start-Stop Assembly cassette.** The plasmid contains the low-copy number origin of replication p15A that allows replication of the shuttle vector in *E. coli*. Left homology arm (LHA) and right homology arm (RHA) mediate integration at the *ssl0410* locus of the *Synechocystis* genome. Between the homology arms is a Level 2 Start-Stop Assembly cassette, which includes a *lacZ*α cassette for blue/white screening, two outward-facing BsaI recognition sites with corresponding A and Z acceptor fusion sites for Level 2 assembly of multiple expression units and two inward-facing BbsI recognition sites with corresponding 1 and 2 donor fusion sites for Level 3 assemblies (see the original Start-Stop Assembly publication for details of these sites[1]). In addition to the Level 2 Start-Stop Assembly cassette, there is a *kanR* gene encoding an aminoglycoside phosphotransferase which confers resistance to kanamycin in both *E. coli* and *Synechocystis*.

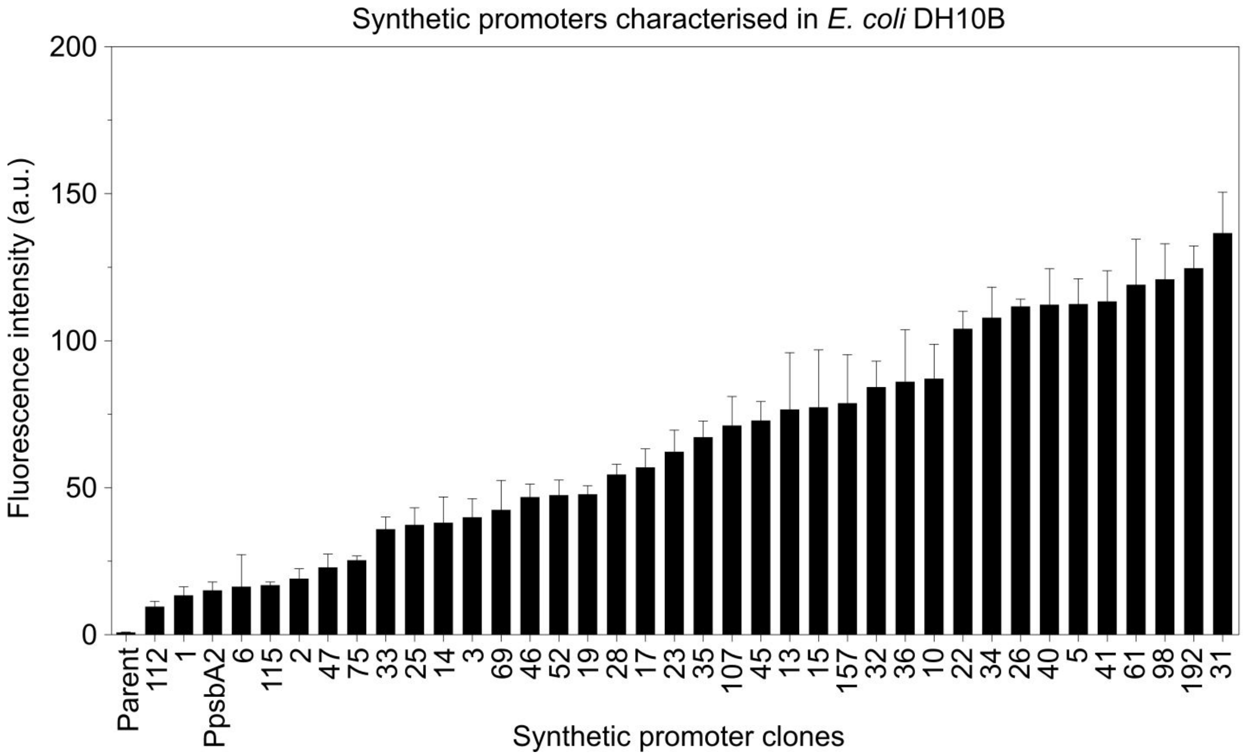


**Figure S2. Synthetic promoter library characterisation in *E. coli* DH10B.** *E. coli* DH10B cells were transformed with plasmids containing the 37 promoter clones derived from the SPL. Transformant colonies were cultured overnight at 37 °C in LB medium supplemented with kanamycin (50 µg $ml^{-1}$), subcultured into fresh LB medium, grown for 4 hours, then the fluorescence of 10,000 cells was measured by flow cytometry. The fluorescence intensities presented represent the fluorescence of cells with wild-type background fluorescence subtracted. 'Parent' plasmid pATM2 was used as template to generate SPL, and uses the native strong promoter Psll1120 to drive expression of EYFP[2]. 'PpsbA2' indicates the plasmid pATM10 used as a positive control, in which the strong native promoter PpsbA2 is used to drive expression of EYFP, using the same insulator and synthetic RBS as the SPL design. The error bars shown represent the standard deviation of five biological replicates.

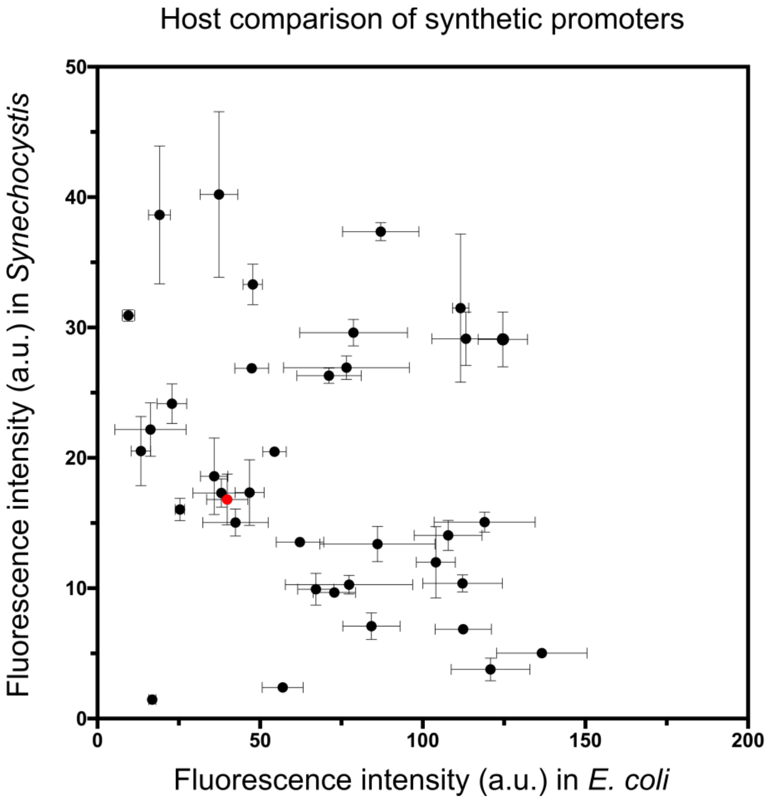


**Figure S3. Comparison of synthetic promoters from the SPL between *E. coli* DH10B and *Synechocystis*.** EYFP fluorescence intensity caused by each synthetic promoter in *E. coli* (as described in Figure S2) was plotted against the EYFP fluorescence intensity caused by the same synthetic promoter in *Synechocystis* (as described in Figure 1b and Figure S8). No correlation in the performance of the synthetic promoters was observed between *E. coli* and *Synechocystis*, $R^2$ = 0.05. SPL promoter clone 3 (red dot) is used in the RBS library design. Error bars represent standard deviations of five biological replicates for *E. coli* and three biological replicates for *Synechocystis*.

| Construct | Sequence |
|---|---|
| pATM2 | TCGGGTAAGTTTATAATATAC**AAAGGAGGT**AGAA |
| RBS library design 1 | NNNNNNNNNNNNNNN**AAAGGAGGT**NNNNN |
| RBS library design 2 | NNNNNNNNNNNNNNN**NAAGGAGGN**NNNNN |
| RBS library design 3 | NNNNNNNNNNNNNNN**NNAGGAGGN**NNNNN |
| RBS library design 4 | NNNNNNNNNNNNNNN**NNNGGAGGN**NNNNN |
| RBS library design 5 | NNNNNNNNNNNNNNN**NNNNGAGGN**NNNNN |
| RBS library design 6 | NNNNNNNNNNNNNNN**NNNNNAGGN**NNNNN |
| RBS library design 7 | NNNNNNNNNNNNNNN**NNNNNNGGN**NNNNN |
| RBS library design 8 | NNNNNNNNNNNNNNN**NNNNNNNGN**NNNNN |
| RBS library design 9 | NNNNNNNNNNNNNNN**NNNNNNNNN**NNNNN |
| pGT77 | AGAGCAGGATTAGAG**GGAGGT**CAGAGA |

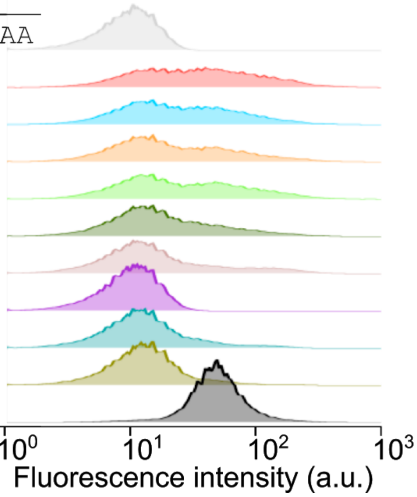


**Figure S4. The distribution of RBS strengths in each of the nine RBS library designs in *E. coli*.** The sequence for each of the library designs shows the different degrees of randomisation ('N' represents an equal mixture of the four bases A, T, C and G) that was introduced in the nine RBS library designs. Flow cytometry histograms show 10,000 *E. coli* cell counts normalised to the maximum (in order to visualise distribution rather than absolute values) for pATM2 as a negative reference, pGT77 as a positive reference and a pool of several hundred transformants for each of the nine library designs.

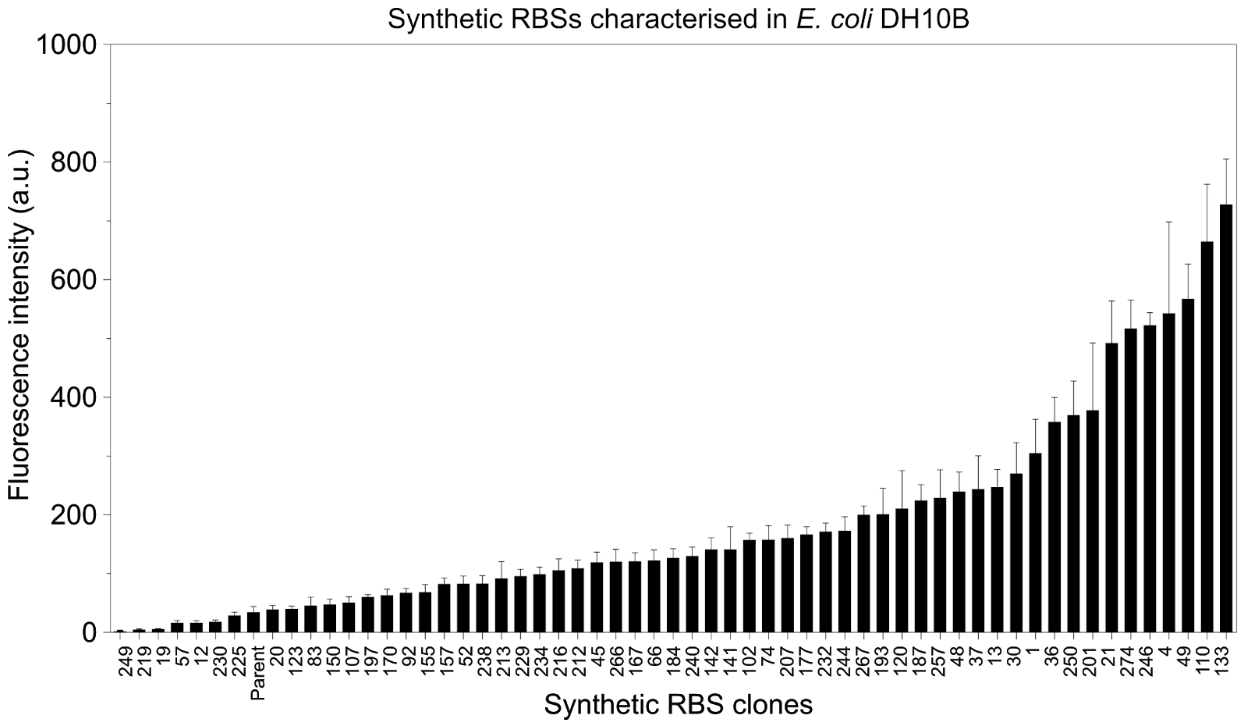


**Figure S5. RBS library characterisation in *E. coli* DH10B.** *E. coli* DH10B cells were transformed with plasmids containing the 58 RBS clones (under promoter SPL clone 3) derived from the RBS library. Transformant colonies were cultured overnight at 37 °C in LB medium supplemented with kanamycin (50 µg $ml^{-1}$), subcultured into fresh LB medium, grown for 4 hours, then the fluorescence of 10,000 cells was measured by flow cytometry. The fluorescence intensities presented represent the fluorescence of cells with wild-type background fluorescence subtracted. ‘Parent’ plasmid pGT77 was used as a template to generate the RBS libraries, and uses the mid-strength promoter SPLc3 which was derived from the SPL. The error bars shown represent the standard deviation of five biological replicates.

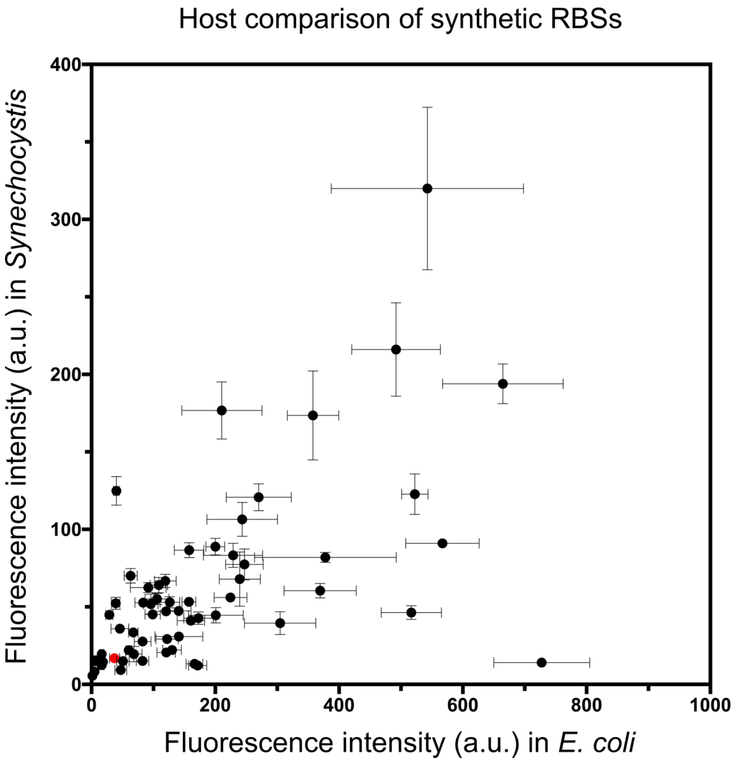


**Figure S6. Comparison of RBSs between *E. coli* DH10B and *Synechocystis* sp. PCC 6803.** EYFP fluorescence intensity caused by each synthetic RBS in *E. coli* (as described in Figure S5) plotted against the EYFP fluorescence intensity caused by the same RBS in *Synechocystis* (as described in Figure 1c). A weak correlation in the performance of the RBSs was observed between *E. coli* and *Synechocystis*, $R^2$ = 0.34, pGT77 (SPL promoter clone 3; red dot) was included as the positive reference containing the promoter used for the RBS library design. Error bars represent standard deviations of five biological replicates for *E. coli* and three biological replicates for *Synechocystis*.

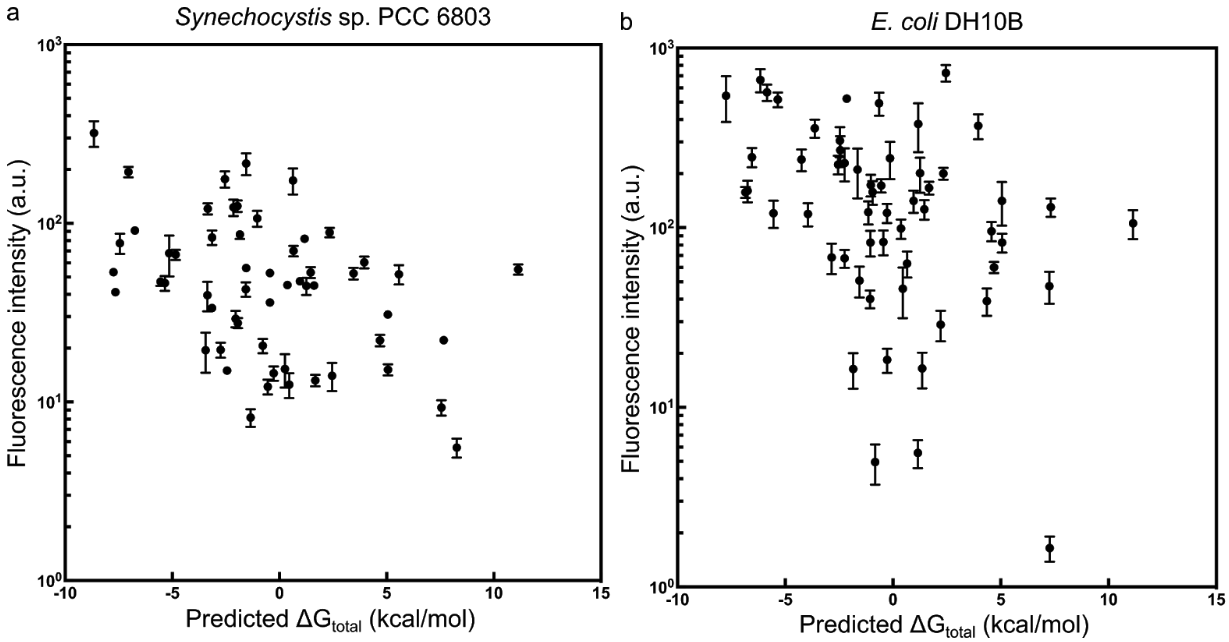


**Figure S7. Comparison of the predicted and measured strength of the synthetic RBSs in *Synechocystis* and *E. coli*.** The log10 EYFP fluorescence caused by the synthetic RBSs compared to their predicted $\Delta G_{tot}$ in (a) *Synechocystis* ($R^2$ = 0.151) and (b) *E. coli* DH10B ($R^2$ = 0.149) calculated using the RBS Calculator v2.0[3]. The error bars shown represent the standard deviation of five biological replicates.

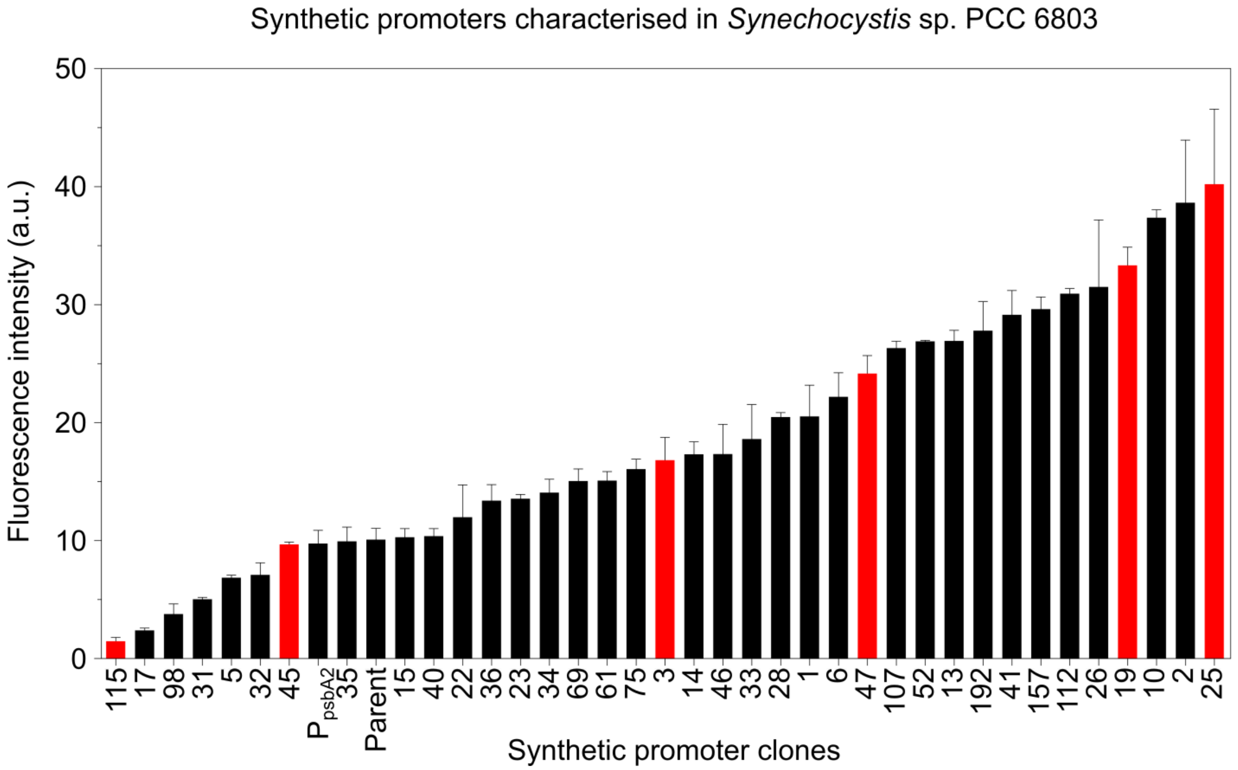


**Figure S8. Synthetic promoter library characterisation in *Synechocystis*.** *Synechocystis* clones with different synthetic promoters derived from the SPL were cultured in BG11 medium in photoautotrophic conditions with constant light and the fluorescence intensity of 10,000 cells was measured in mid-linear phase of growth using flow cytometry. Fluorescence intensities presented represent the fluorescence of cells with background fluorescence subtracted. 'Parent' represents a transformant of the plasmid pATM2, from which the synthetic promoter library was derived. 'PpsbA2' indicates the plasmid pATM10 used as a positive control, the plasmid uses the strong native promoter PpsbA2 to drive expression of EYFP, using the same insulator and synthetic RBS as the SPL design. The red bars represent the subset of six chosen promoters. The error bars shown represent the standard deviation of three biological replicates.

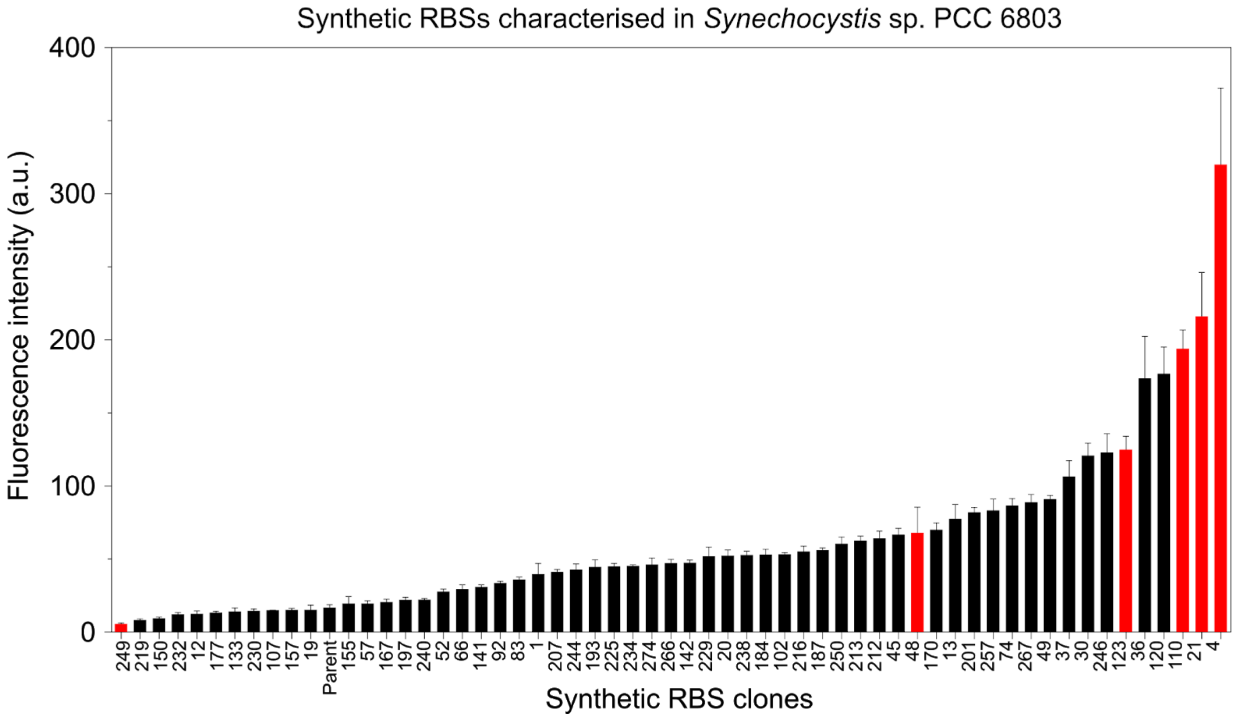


**Figure S9. RBS library characterisation in *Synechocystis* sp. PCC 6803.** *Synechocystis* clones with different synthetic RBSs derived from the RBS library were cultured in BG11 medium in photoautotrophic conditions with constant light and the fluorescence intensity of 10,000 cells was measured in mid-linear phase of growth using flow cytometry. Fluorescence intensities presented represent the fluorescence of cells with background fluorescence subtracted. 'Parent' indicates a transformant of the plasmid pGT77, from which the RBS library was derived. The red bars represent the subset of six chosen RBSs. The error bars shown represent the standard deviation of three biological replicates.

**Design and assembly of rational pathway designs (pGT432, pGT433, pGT434 and pGT435)**

**Design**

| | | |
|---|---|---|
| High expression design pGT432 | SPLc19 | RBSc21 |
| Medium expression design pGT433 | SPLc3 | RBSc123 |
| Low burden design pGT434 | SPLc19 | RBSc48 |
| High burden design pGT435 | SPLc45 | RBSc21 |

**Assembly**

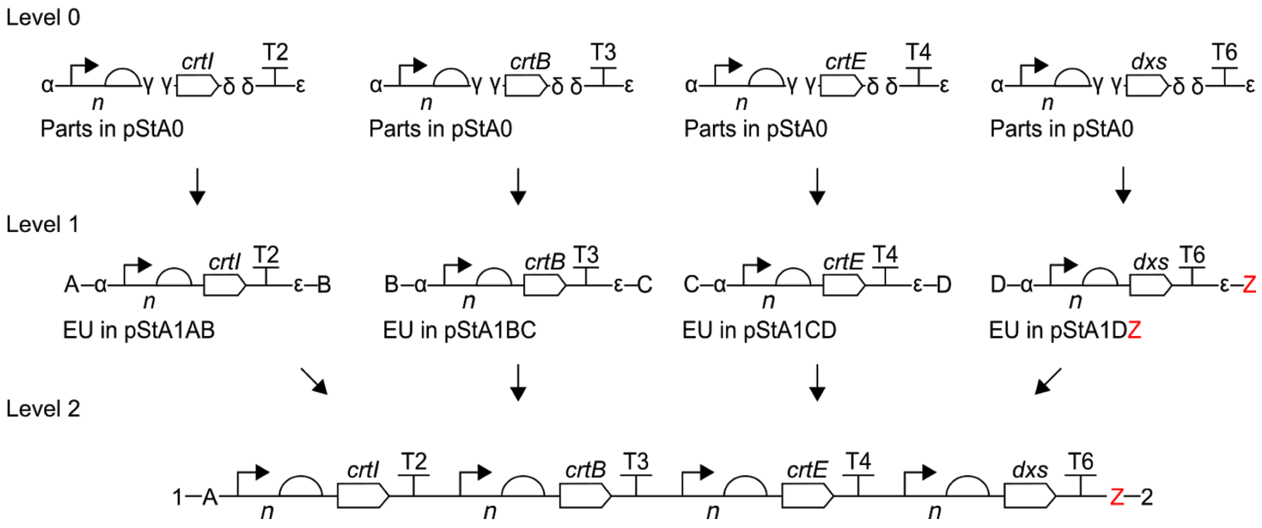


**Figure S10. Design and assembly of four individual pathway-encoding constructs.** Four pathway design strategies were used: high expression (pGT432), medium expression (pGT433), low burden (pGT434) and high burden (pGT435). Each construct was individually assembled using Start-Stop Assembly in the destination vector pGT270. pGT434 was not cloneable in *E. coli* in three independent attempts. '*n*' shows the uncertain representation of each promoter and RBS at each position following assembly. T2 = terminator L3S2P21, T3 = terminator ECK120033737, T4 = terminator ECK120019600 and T6 = ECK120015170. Assembled insert size = 5818 bp. Total size of plasmids including assembled insert = 10,164 bp.

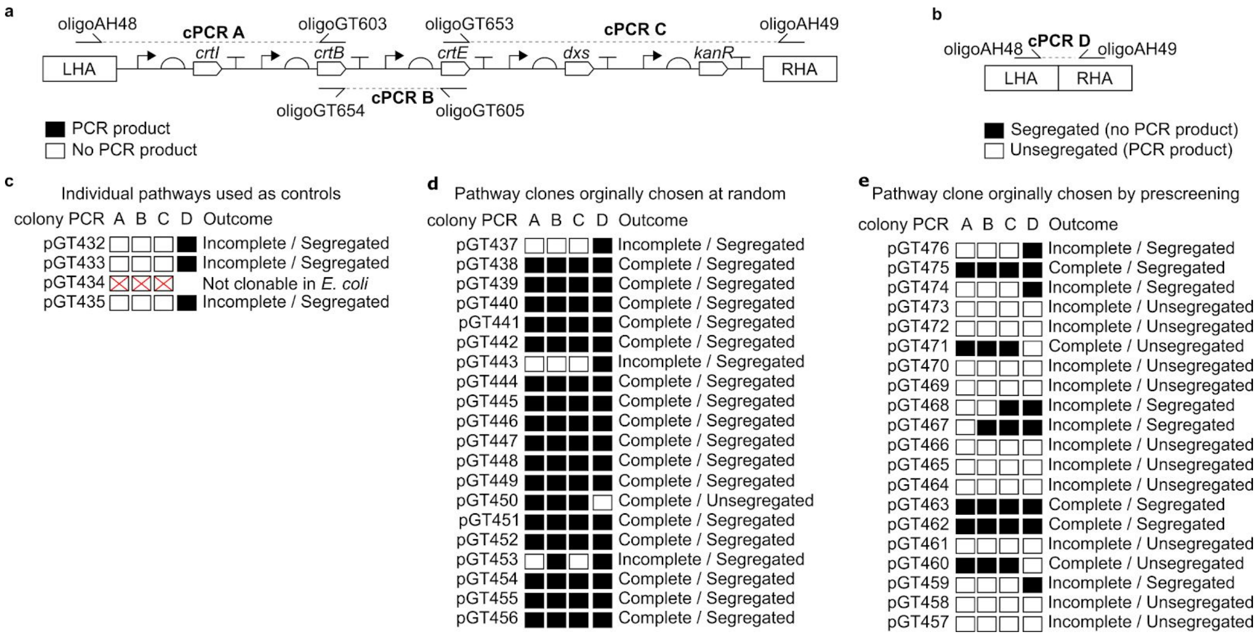


**Figure S11. PCR screening for genetic stability of pathway constructs in *Synechocystis*.** (a) To confirm whether each pathway construct was complete or incomplete, three separate colony PCR reactions (A, B and C) were used, because complete pathways were too large to amplify as one fragment. (b) To assess segregation colony PCR D was used, which tested for the presence of a wild-type copy of the integration locus. The presence of a wild-type copy of the locus indicates that the introduced pathway had not fully segregated. The absence of the wild-type amplicon suggests that the pathway construct was present in every copy of the chromosome. Complete pathways were too large to amplify therefore this reaction only screened for the presence or absence of the wild-type locus. (c) The outcome of the four colony PCR reactions for the four rational pathway designs. (d) The outcome of the four colony PCR reactions for the 20 pathway variants originally chosen at random. (e) The outcome of the four colony PCR reactions for the 20 pathway variants originally chosen by prescreening in *E. coli*. Black boxes represent the desired outcome, white boxes represent the unwanted outcome. In the case of PCR reactions A, B and C, black boxes represent the presence of that pathway amplicon, whereas white boxes represent its absence. In the case of PCR reaction D, a black box represents a segregated pathway construct, whereas a white box represents an unsegregated pathway construct.

Assembly of pGT547

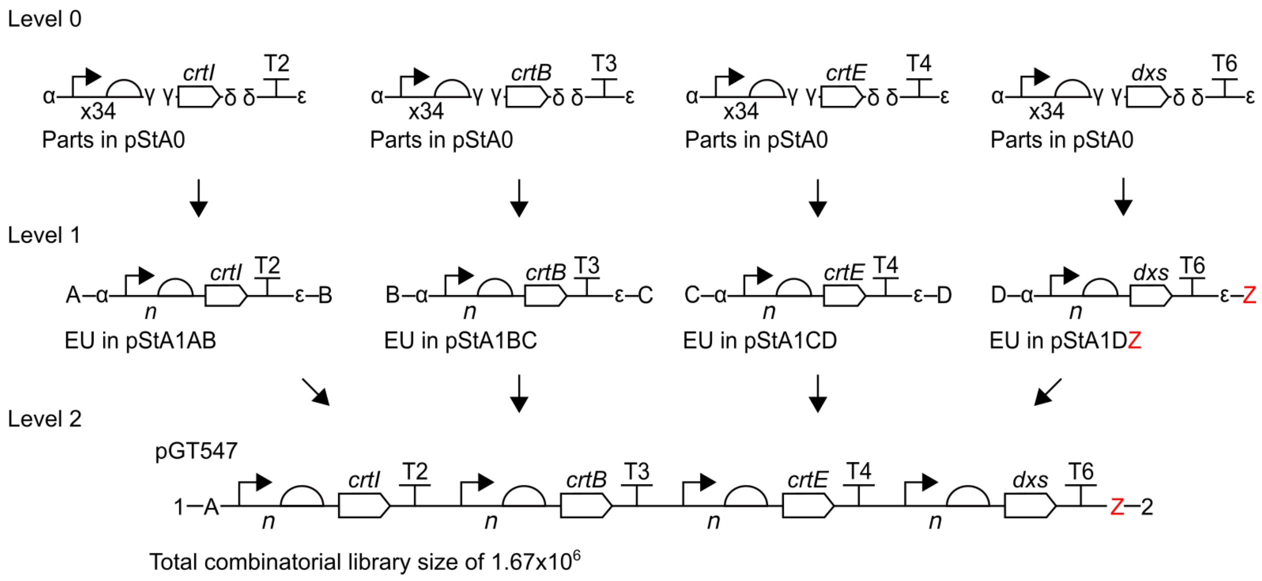


**Figure S12. Combinatorial assembly of lycopene pathway library pGT547.** A library of lycopene pathway construct variants was assembled as shown using Start-Stop Assembly. The destination vector was pGT270. An equimolar mixture of the 34 promoter-RBS composite parts labelled as 'x34' was used in Level 1 assemblies. '*n*' shows the uncertain representation of each promoter and RBS at each position following assembly. The maximum library size was $34^4$ = $1.3 \times 10^6$. T2 = terminator L3S2P21, T3 = terminator ECK120033737, T4 = terminator ECK120019600 and T6 = ECK120015170. Assembled insert size = 5818 bp. Total size of plasmids including assembled insert = 10,164 bp.

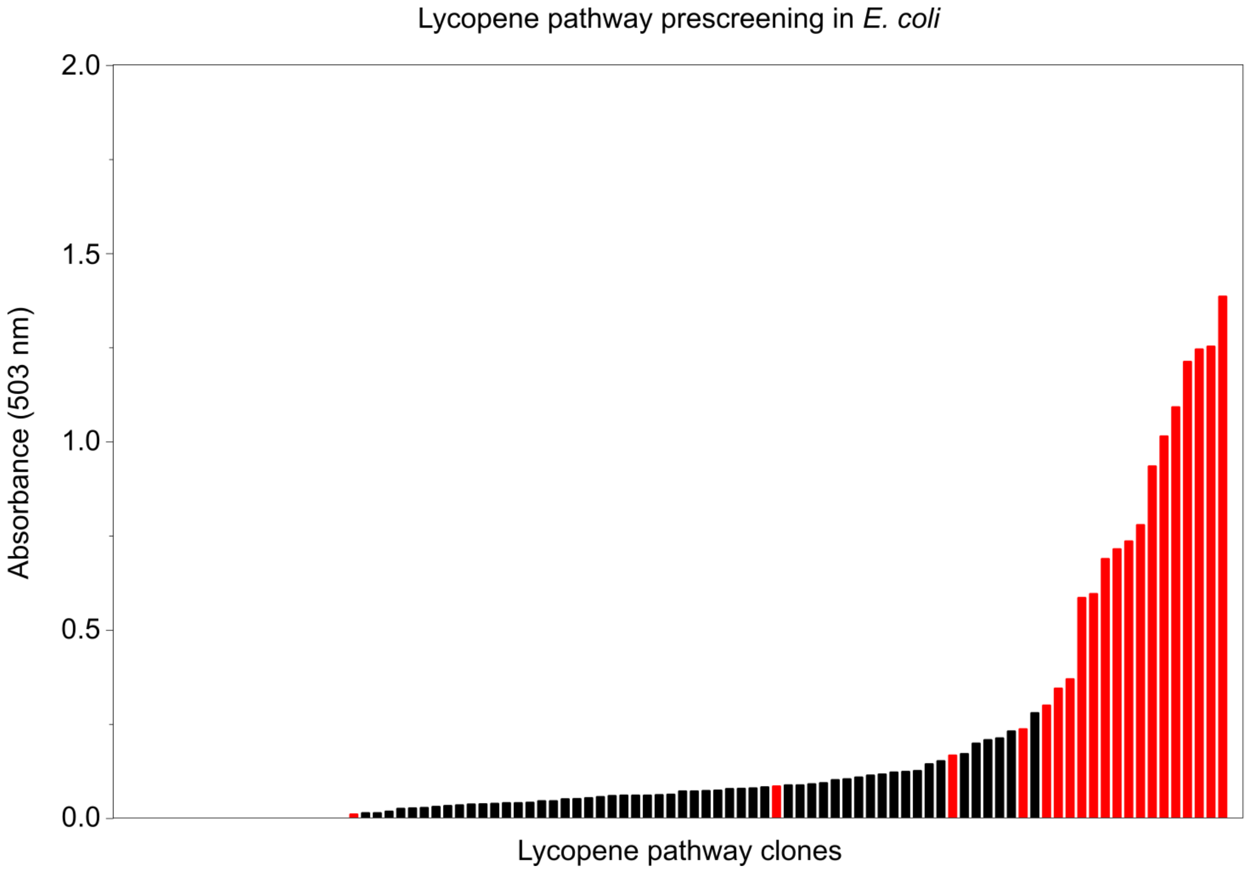


**Figure S13. Prescreening of construct variants for lycopene production in *E. coli* by absorbance**. As described in the main text, 96 *E. coli* transformant clones were picked at random from the combinatorial lycopene overproduction pathway library and screened for lycopene production after 48 h of growth by spectrophotometry (as described in Methods). The absorbance at 503 nm of each clone is presented with background absorbance subtracted. Of these 96 clones, twenty (shown in red) were chosen for further study, of which ten were the strongest lycopene producers and the other ten were evenly-distributed throughout the observed range of lycopene concentrations. Order of pathway clones from least to most lycopene produced, left to right: clone 34, 67, 93, 10, 70, 86, 5, 14, 46, 51, 74, 89, 37, 22, 41, 49, 61, 91, 78, 94, 58, 44, 82, 39, 64, 54, 50, 27, 15, 53, 19, 32, 38, 56, 59, 33, 43, 85, 52, 3, 88, 30, 28, 25, 47, 48, 57, 62, 9, 83, 8, 11, 68, 18, 80, 26, 2, 31, 65, 71, 84, 40, 36, 96, 6, 73, 55, 60, 1, 81, 66, 63, 35, 72, 13, 42, 16, 23, 12, 45, 7, 21, 79, 87, 17, 75, 4, 24, 90, 95, 77, 92, 20, 69, 76.

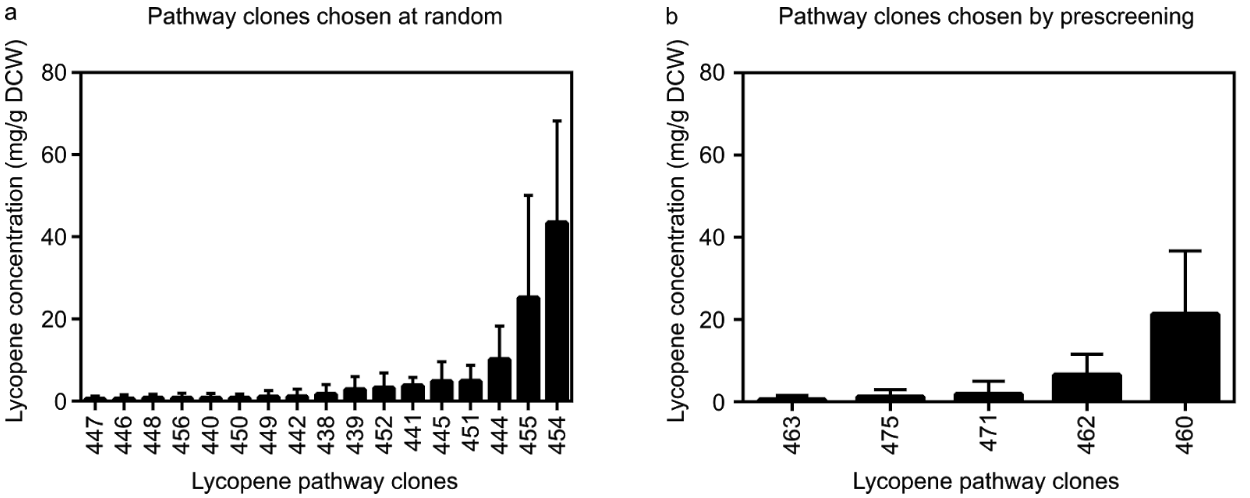


**Figure S14. Lycopene concentrations in *E. coli* produced by pathway construct variants which were stable in *Synechocystis*.** *E. coli* DH10B cells were transformed with the lycopene overproduction pathway construct variants chosen (a) at random and (b) after prescreening in *E. coli*. Only those construct variants which had proven genetically stable in *Synechocystis* are shown. Transformant clones were grown for 24 hours at 30 °C in LB medium supplemented with kanamycin (50 µg $ml^{-1}$), subcultured into fresh LB medium, grown for 48 hours, then lycopene concentration was determined. The error bars shown represent the standard deviation of three independent biological replicates.

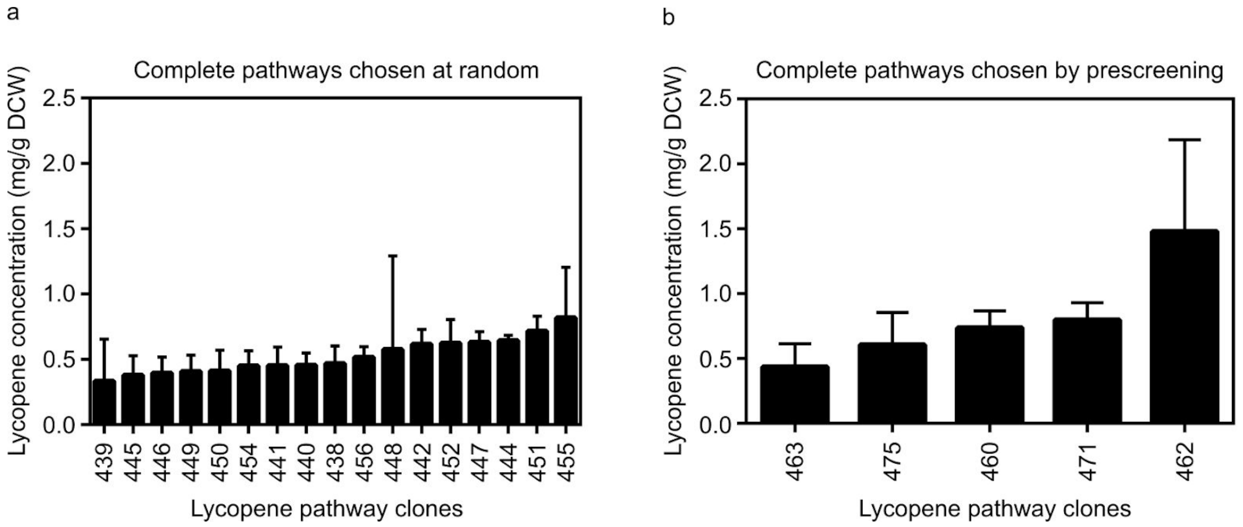


**Figure S15. Lycopene concentrations in *Synechocystis* produced by pathway construct variants, grouped by strategy originally used to identify variants.** *Synechocystis* cells containing complete pathway constructs (a) chosen at random in *E. coli* and (b) chosen by prescreening in *E. coli* were grown in BG11 medium supplemented with kanamycin (30 μg $ml^{-1}$), in photoautotrophic conditions and constant light for 2 days, subcultured into fresh BG11 medium, grown for two weeks, then lycopene concentration was determined. The error bars shown represent the standard deviation of three independent biological replicates. Variant names are shown as numbers only, omitting the initial ‘pGT’.

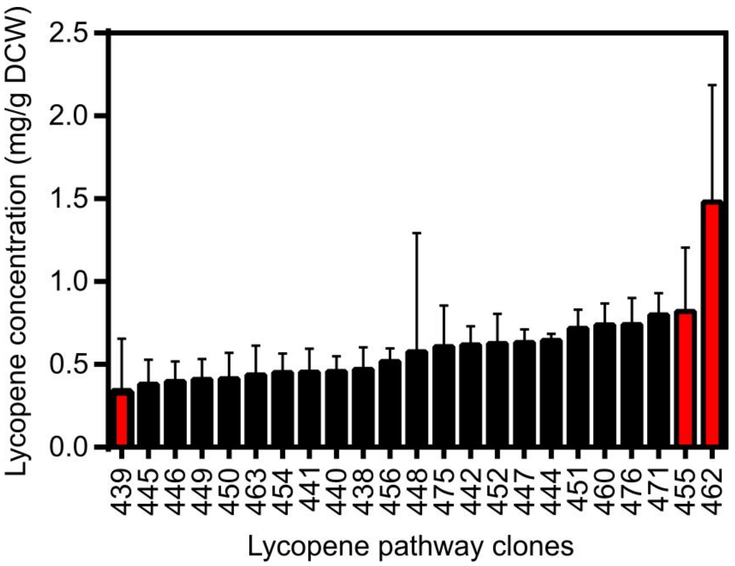


**Figure S16. Lycopene concentrations in *Synechocystis* produced by pathway construct variants, showing three variants chosen for further study.** Lycopene concentrations of the 17 pathway variants chosen at random (Figure 4) and the six pathway variants chosen by prescreening in *E. coli* (Figure 4) characterised in *Synechocystis*. Three variants chosen for further characterisation are shown in red: a low producing strain pGT439, a medium producing strain pGT455, and a high producing strain pGT462 . Variant names are shown as numbers only, omitting the initial ‘pGT’.

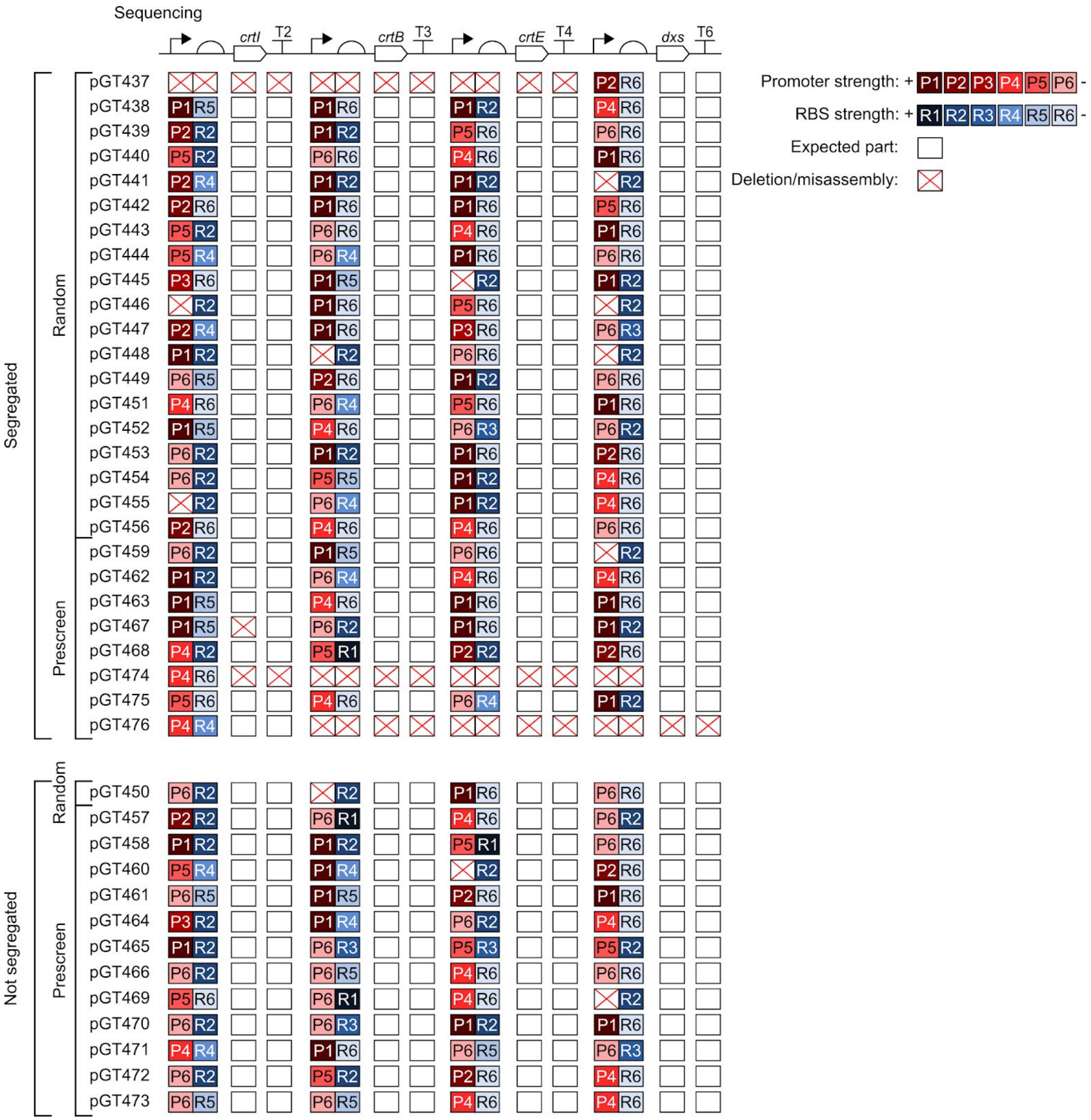


**Figure S17. Sequencing results of lycopene pathway variants grouped first by pathway segregation, then by strategy used to identify variants.** All 40 pathway variants (20 pathway variants chosen at random and 20 pathway variants chosen by prescreening) were sequenced. Promoters, in descending order of strength as characterised using EYFP: P1 = SPLc25, P2 = SPLc19, P3 = SPLc47, P4 = SPLc3, P5 = SPLc45 and P6 = SPLc17. RBSs, in descending order of strength as characterised using EYFP: R1 = RBSc4, R2 = RBSc21, R3 = RBSc110, R4 = RBSc123, R5 = RBSc48 and R6 = RBSc249. In the sequencing results the white box indicates presence of the expected part and the red cross represents a deletion or misassembly.

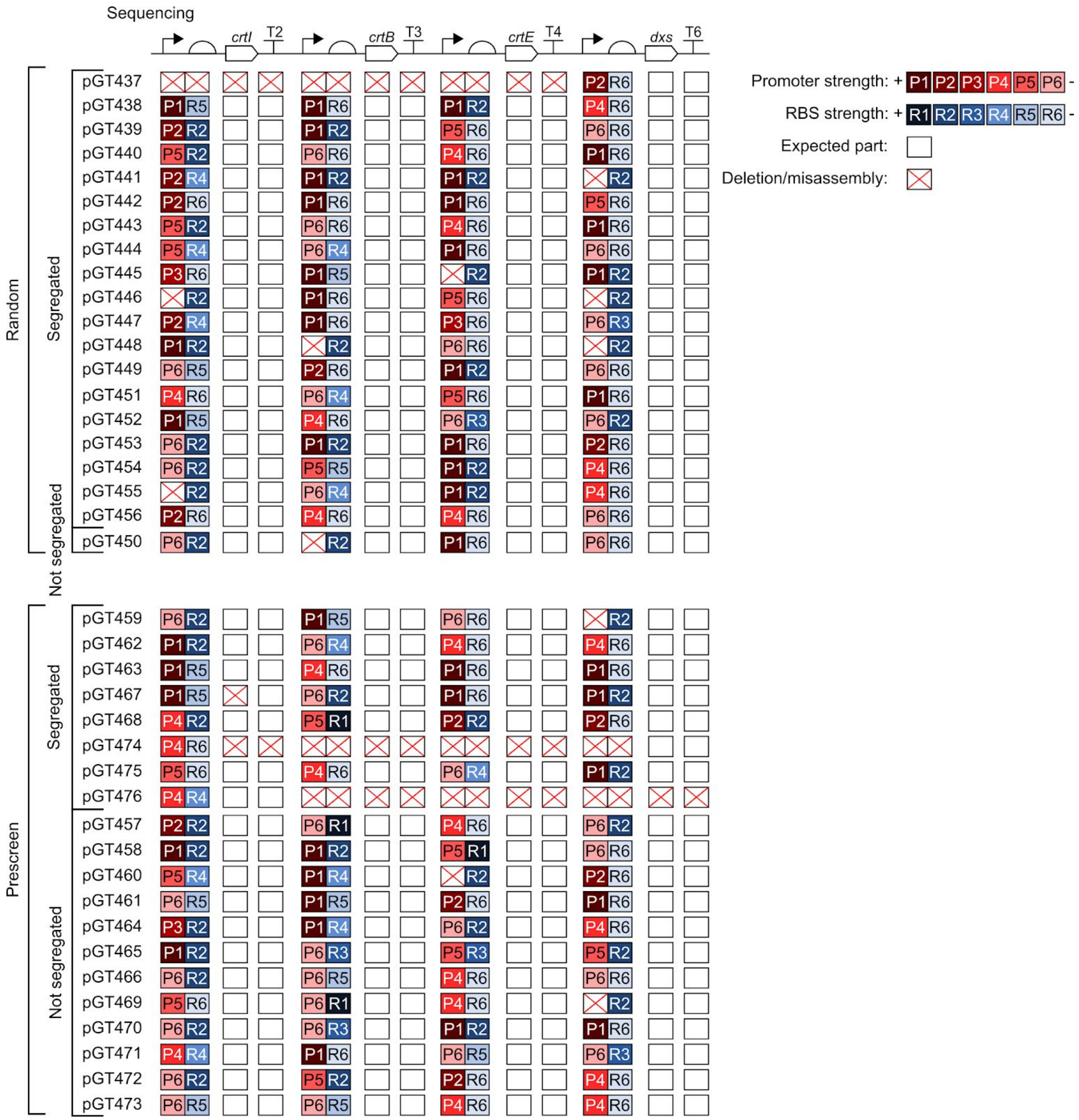


**Figure S18. Sequencing results of lycopene pathway variants grouped first by strategy used to identify variants, then by pathway segregation.** All 40 pathway variants (20 pathway variants chosen at random and 20 pathway variants chosen by prescreening) were sequenced. Promoters, in descending order of strength as characterised using EYFP: P1 = SPLc25, P2 = SPLc19, P3 = SPLc47, P4 = SPLc3, P5 = SPLc45 and P6 = SPLc17. RBSs, in descending order of strength as characterised using EYFP: R1 = RBSc4, R2 = RBSc21, R3 = RBSc110, R4 = RBSc123, R5 = RBSc48 and R6 = RBSc249. In the sequencing results the white box indicates presence of the expected part and the red cross represents a deletion or misassembly.

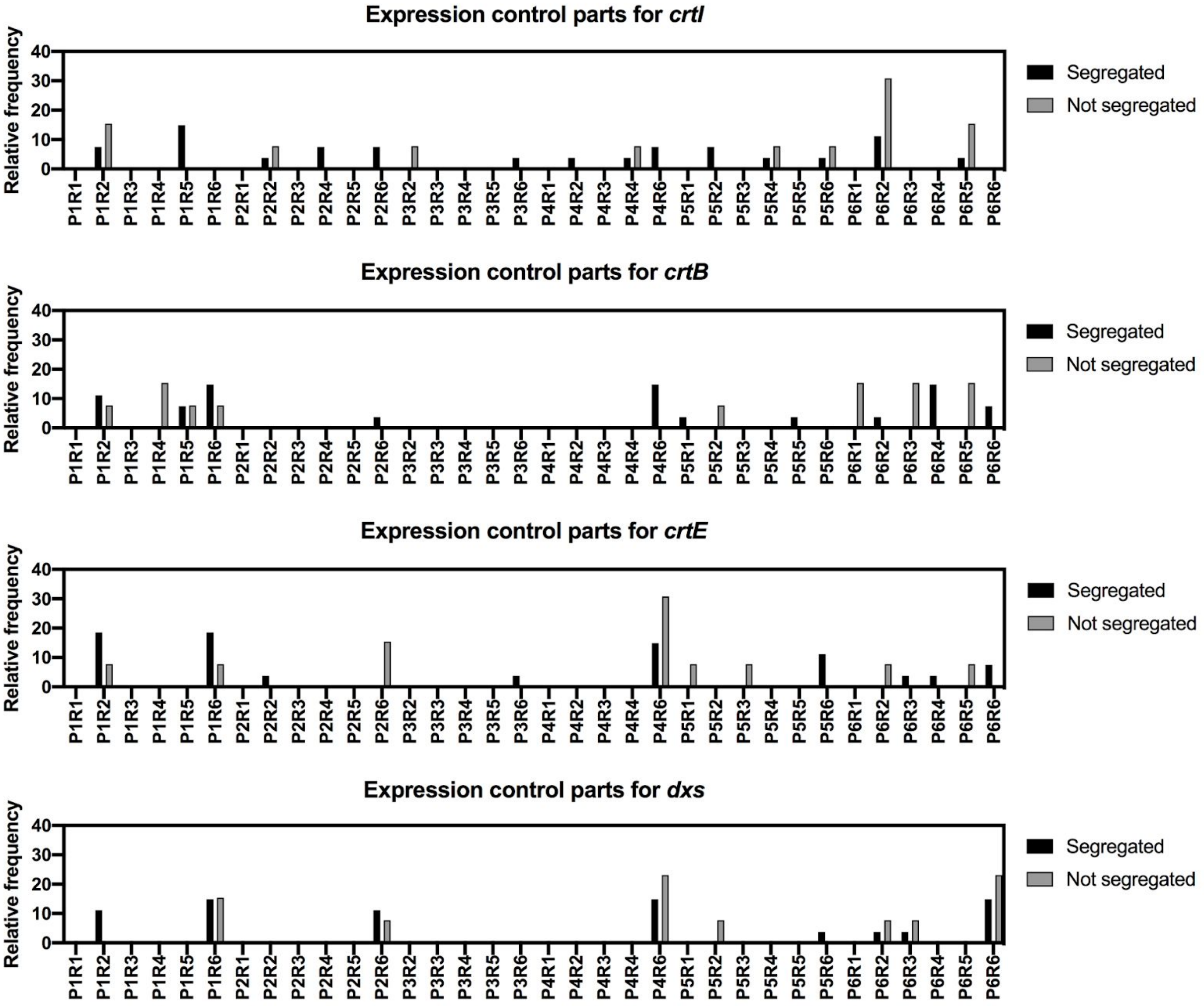


**Figure S19. Frequencies of promoter and RBS combinations in pathway variants.** The relative frequencies of each promoter and RBS combination found in segregated and unsegregated pathway variants. Segregated pathway variants are displayed as black bars. Non-segregated pathway variants are displayed as grey bars. Promoters, in descending order of strength as characterised using EYFP: P1 = SPLc25, P2 = SPLc19, P3 = SPLc47, P4 = SPLc3, P5 = SPLc45 and P6 = SPLc17. RBSs, in descending order of strength as characterised using EYFP: R1 = RBSc4, R2 = RBSc21, R3 = RBSc110, R4 = RBSc123, R5 = RBSc48 and R6 = RBSc249.

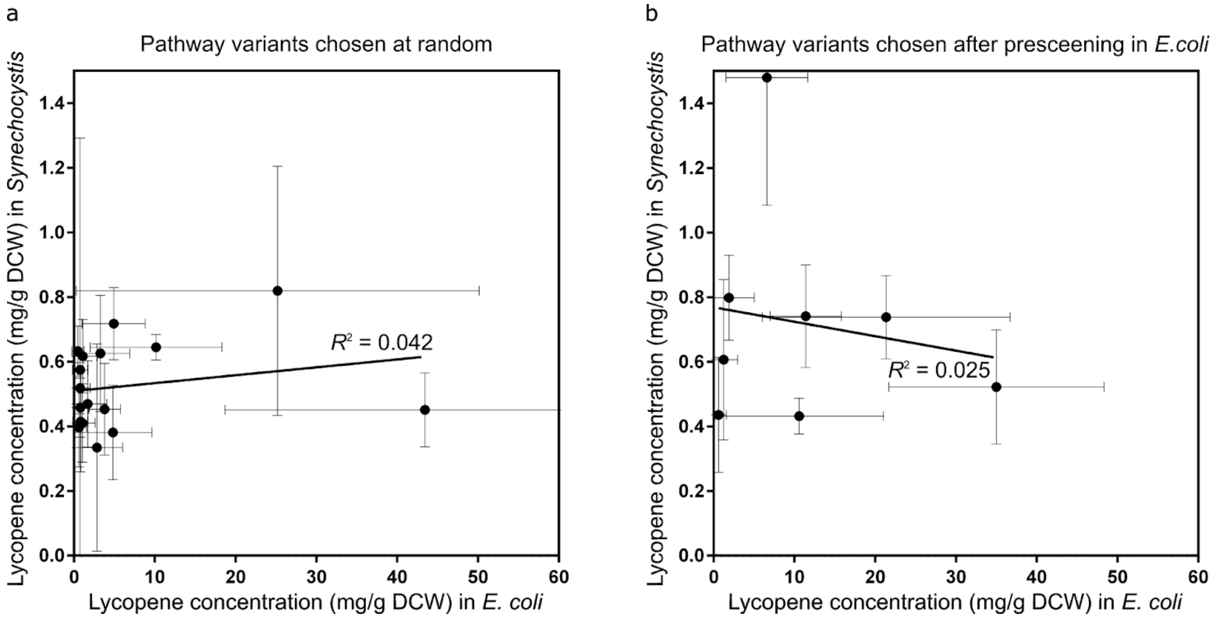


**Figure S20. Comparison of lycopene concentrations produced by pathway construct variants in *E. coli* and *Synechocystis*.** (a) Lycopene concentrations in *E. coli* (Figure S14a) compared against those in *Synechocystis* (Figure S17a) for the pathway variants chosen at random. (b) Lycopene concentrations in *E. coli* (Figure S14b) compared against those in *Synechocystis* (Figure S17b) for the pathway variants chosen by prescreening in *E. coli*. Error bars represent standard deviations of three independent biological replicates for *E. coli* and *Synechocystis*.

**Table S1. Sequences of synthetic promoters from the synthetic promoter library.** -35 and -10 regions are shown in **bold**.

| Synthetic promoter | Sequence (5' - 3')<br>-35 region -10 region |
|---|---|
| 1 | CATCGCCCGA**TTGACA**CCTCGCGCCACGCCTGA**TATAAT**GGGGGG |
| 2 | TCAAGGGAAG**TTGACA**CGACACCGCTAATTTAT**TATAAT**GAAGAG |
| 3 | CCAGATGACC**TTGACA**AGTAACTGCCCCAACAG**TATAAT**ATACCT |
| 5 | CAGTGTGGCG**TTGACA**TTCAGTCATGCACCCCG**TATAAT**TGGCGA |
| 6 | GCGGGGTTGC**TTGACA**GGATTTACTCGGCAGGG**TATAAT**GGGTAC |
| 10 | CGCAGCGGGC**TTGACA**CATCGGCCCAAACTACA**TATAAT**GTGTAT |
| 13 | CCCGCAGCTT**TTGACA**CACCGCCACTGACAGTC**TATAAT**TTGGAT |
| 14 | TAGTATAAGA**TTGACA**TAATGTAGTCCGTGGAG**TATAAT**GTAATT |
| 15 | CACCAGAGGA**TTGACA**GTCCTAGCAGACACGTG**TATAAT**AGTCCC |
| 17 | CCCCTAAGCT**TTGACA**CGCAGAGAACACAACCC**TATAAT**GATTCG |
| 19 | GCGGCTCGTA**TTGACA**ATTTAACTAAATATCAG**TATAAT**TTAGTT |
| 22 | ATAGGCACCG**TTGACA**TACTAGCCAGAAGAAAA**TATAAT**TGGGGG |
| 23 | CGGGTGGCAA**TTGACA**TGGCACGAGTCCGTCTA**TATAAT**AACAGA |
| 25 | CCACGAAAAA**TTGACA**AGCACTCTGCGCTCTGA**TATAAT**TAAGGT |
| 26 | CCCCCACCCC**TTGACA**GCCCGTGGCTCGATGCA**TATAAT**GAATGT |
| 28 | CCCACATAAC**TTGACA**GAAGTCCGCAGCGACTT**TATAAT**ACGGGG |
| 31 | TTTTATTATA**TTGACA**TCTTTCATACACCTGCA**TATAAT**TCGAGT |
| 32 | CGGTCGGCCT**TTGACA**GCGGTTGATTAAGAGAC**TATAAT**GTCTAT |
| 33 | CTCGCGGCAA**TTGACA**AAATTGGATCTCGATAT**TATAAT**GGGGGT |
| 34 | CCGACATCCT**TTGACA**CGAACTCGCTGGCCCGG**TATAAT**TCTTTA |
| 35 | ACACGCCGCC**TTGACA**TCACTTTACGGCACCCA**TATAAT**GGGGGG |
| 36 | GCCCCCCGCG**TTGACA**CCCGCCCCGTACTCGCC**TATAAT**TAAGGG |
| 40 | CCAAATTATA**TTGACA**TTACCGGGGTCGTGAAC**TATAAT**TCTTTC |
| 41 | TGCCCCCCCC**TTGACA**ACCGCTGTCCCGCATCC**TATAAT**TGTTCA |
| 45 | CTCGCCACCT**TTGACA**CAGCGGCCCGCTAACTA**TATAAT**AAACGT |
| 46 | AGCGCGGACC**TTGACA**CTCCTAGTGACTTACAC**TATAAT**AGCTGG |
| 47 | GTGACGTCTA**TTGACA**GCCAGGAGTTACCGAGA**TATAAT**GGAATA |
| 52 | CCCCCTCCCC**TTGACA**CCCTCCCATCCCGATGT**TATAAT**GTGTTT |
| 61 | TCTAGCCAGA**TTGACA**CTGACATTAAGTACACT**TATAAT**CGGAGG |
| 69 | GTCGCCCCCA**TTGACA**AATGTAAGTCGCGGAGG**TATAAT**AGGTGC |
| 75 | CTCACGGACC**TTGACA**GACGGGGGCCCCCAGG**TATAAT**GAGGGG |

| | |
|---|---|
| 98 | ACCATCTAGT**TTGACA**TCGAAAAGCCAACAACG**TATAAT**CAGCTT |
| 107 | CGCGCTACCA**TTGACA**TCCCGTTCCCGCCGCTG**TATAAT**TGATTT |
| 112 | CGAAAACACA**TTGACA**CCCTTTTATAATCATGA**TATAAT**TGGATC |
| 115 | CCGGACCCCC**TTGACA**CTACGGCCCGCTATGGA**TATAAT**TCACCG |
| 157 | CACCGGGCCT**TTGACA**ATACGTATCAAGTACCT**TATAAT**AAGGGG |
| 192 | CATTCCGTGC**TTGACA**CATTATTAAACTATGAA**TATAAT**GTGTAA |

**Table S2. Sequences of synthetic RBSs from the RBS library.**

| RBS library design | Synthetic RBS | Sequence (5' - 3') |
|---|---|---|
| RBS library design 1 | 1 | CTCGATTTCGCGAACAAAGGAGGTGAGGT |
| | 4 | TATGTCAGTAAAAAAAAAGGAGGTTCGTT |
| | 12 | TAAAGTTAAGCGTCCAAAGGAGGTTTAAC |
| | 13 | GCGGATAGTGAGGAGAAAGGAGGTGGGGG |
| | 19 | ACAAACTCCTCACGGAAAGGAGGTACGGA |
| | 20 | GCAAATTCTCGTGGGAAAGGAGGTCGCGG |
| | 21 | TACGAAAGATTTACGAAAGGAGGTTGGTT |
| | 30 | ATTTCGTATGAAGGTAAAGGAGGTTGGCA |
| | 36 | CTCTAGGGTTTTCTAAAGGAGGTTGAAT |
| | 37 | AGTTGACCGTAACGCAAAGGAGGTAGAGC |
| | 45 | TCGGCGGCGTAAACTAAAGGAGGTGGAAG |
| | 48 | AGTGTGTTCCAAGAGAAAGGAGGTATCTG |
| | 49 | AATAAAAGCCAGTTTAAAGGAGGTCTTTG |
| | 52 | ATTAGACGACGATCCAAAGGAGGTCCCAG |
| | 57 | TGGACTCCTACGAGGAAAGGAGGTGGTA |
| | 66 | CTTATGGGTGTTTGAAAGGAGGTGGACG |
| | 74 | AGTACTAACGGGTTTAAAGGAGGTCGCGG |
| | 83 | GCGGATAGTGAGGAGAAAGGAGGTGGGGG |
| | 92 | AAACCGAACAGGCCAAAGGAGGTCAGCT |
| | 102 | TTTAGTATTGCTACTAAAGGAGGTTGCAC |
| | 107 | TTGGGGCGATTTGGGAAAGGAGGTGCGGG |
| | 110 | TAGGTATATGCAATCAAAGGAGGTATATG |
| | 120 | ATACTGCATCAATGAAAGGAGGTAGGGG |
| | 123 | TGTGTAGGTCCCACCAAAGGAGGTCTGTA |
| RBS library design 4 | 133 | TTACTATAATAACGTTTCGGAGGGAATAT |
| | 141 | TTCAAATAAGCATTGATCGGAGGGTTTAC |
| | 142 | GGGATATAACGAAGAGGCGGAGGAGTGCG |
| | 150 | TGTTAAGAGGGGGACTGTGAGGGCTGAG |
| | 155 | TTACCGGTCAAGGCAAGAGGAGGCTAGTC |
| | 157 | TATTATGTGATAATTAGAGGCGTGCA |
| | 167 | GTATATGGTGGAGGAGGTAGCGT |
| | 170 | CTGATTTAGGGGCCATTTTGAGGAATATG |
| | 177 | GTAGGTTGGATATCTGGAGGTGAAGC |

| | | |
|---|---|---|
| RBS library design 4 continued | 184 | CTGCGTGACAACAGATGTCGAGGTGAAAG |
| | 187 | CTCTATTGTGTACTATGAGGAGGCAGCGT |
| | 193 | GTTGCGTAACAATAGACGGAGGATGCTT |
| | 197 | CTGGTATGGTTCAACGAGGATCGTT |
| | 201 | CTATCGGTTATGTGCTAGCGAGGTTTAAT |
| | 207 | GTGCAGACGATGAGAAGGGAGGAAAAGA |
| | 212 | TTTCGAGGAGGATAGCG |
| | 213 | TTTCGAGGAGGATAGCG |
| | 216 | TTTCGAGGAGGATAGCG |
| | 219 | GAGGGTATAGTGCGTAACGGAGGGGTACG |
| | 225 | GTAACGGGGAGTTTAAGTAGAGGGGGTGT |
| | 229 | CGGTGTCTTTATTCTAATTGAGGGGGCTT |
| | 230 | TTGAAAGGGACCCCACAGTGAGGTAGAGC |
| | 232 | TAGGCGTAAGGATCTATCGAGGAAGTGT |
| | 234 | TAATGGTCAGCGTGTGCGGAGGAGCGTG |
| | 238 | TACGGGTTAGTTTGTTGTGGAGGTTACTT |
| | 240 | TCAATTGCTTTGGCGGTGAGAGGCGTGTT |
| | 244 | TGGACATTCAAATATTGAAGAGGGGTGTA |
| | 246 | CCCTATTAAAAACAGACGGAGGCAGAAT |
| RBS library design 9 | 249 | GGCGCGAGTCAACGTTCCTGAGGAGTGGC |
| | 250 | TCTTTGAGACTTTATTGTGGAGGTTTGGC |
| | 257 | CGGAGCAATCAACGAAAGGGGGAAGAGGG |
| | 266 | CGGAGCAATCAACGAAAGGGGGAAGAGGG |
| | 267 | GTTTCTTAGGTGTTGGAGGTTGGTCTA |
| | 274 | AGAGCATTAGGTGATAGGAGGTAAGTA |

**Table S3. Oligonucleotides used in the study.** 5’-phosphorylated oligonucleotides denoted /5Phos/. Fw denotes forward primer. Rv denotes reverse primer.

| Oligonucleotide | Sequence (5’ – 3’) | Comments |
|---|---|---|
| oligoATM3 | /5phos/NNNNNNNNNNNNNNNNNTGTCAAN NNNNNNNNNCTGCAGTCTCCTTTAAATCGG | Rv primer for generating SPL library in pATM2 |
| oligoATM5 | /5phos/TATAATNNNNNNCCGGCTTCGGCG GTAAGGTAAGAGCAGGATTAGAGGGAGGTCA GAGAATGGTGAGCAAGGGCGAGG | Fw primer for generating SPL library in pATM2 |
| oligoATM51 | /5phos/AGGTATATTATACTGTTGGGGCAG | Rv primer for generating RBS library in pATM2 |
| oligoATM52 | NNNNNNNNNNNNNNNAAAGGAGGTNNNNNAT GGTGAGCAAGGGCGAGG | Fw primer for generating RBS library design 1 in pATM2 |
| oligoATM53 | NNNNNNNNNNNNNNNNAAGGAGGNNNNNNAT GGTGAGCAAGGGCGAGG | Fw primer for generating RBS library design 2 in pATM2 |
| oligoATM54 | NNNNNNNNNNNNNNNNNAGGAGGNNNNNNAT GGTGAGCAAGGGCGAGG | Fw primer for generating RBS library design 3 in pATM2 |
| oligoATM55 | NNNNNNNNNNNNNNNNNGGAGGNNNNNNNAT GGTGAGCAAGGGCGAGG | Fw primer for generating RBS library design 4 in pATM2 |
| oligoATM56 | NNNNNNNNNNNNNNNNNNGAGGNNNNNNNAT GGTGAGCAAGGGCGAGG | Fw primer for generating RBS library design 5 in pATM2 |
| oligoATM57 | NNNNNNNNNNNNNNNNNNNAGGNNNNNNNAT GGTGAGCAAGGGCGAGG | Fw primer for generating RBS library design 6 in pATM2 |
| oligoATM58 | NNNNNNNNNNNNNNNNNNNNGGNNNNNNNAT GGTGAGCAAGGGCGAGG | Fw primer for generating RBS library design 7 in pATM2 |
| oligoATM59 | NNNNNNNNNNNNNNNNNNNNNGNNNNNNNAT GGTGAGCAAGGGCGAGG | Fw primer for generating RBS library design 8 in pATM2 |
| oligoATM60 | NNNNNNNNNNNNNNNNNNNNNNNNNNNNNAT GGTGAGCAAGGGCGAGG | Fw primer for generating RBS library design 9 in pATM2 |
| oligoGT234 | GGGGAAACGCCTGGTATCT | pStA0 Fw sequencing primer |

| | | |
|---|---|---|
| oligoGT235 | AGCAAAAACAGGAAGGCAAA | pStA0 Rv sequencing primer |
| oligoGT339 | GTTGAGGACCCGGCTAGG | pStA1 Fw sequencing primer |
| oligoGT340 | TGTGACGGAAGATCACTTCG | pStA1 Rv sequencing primer |
| oligoGT373 | TACTGGGTCTCTCTCCGGCATTGTCTTCcat aactgcagtctcctttaaatcggtttc | Fw primer for constructing pGT270 |
| oligoGT374 | /5Phos/ctccaggatccaaagccacg | Rv primer for constructing pGT270 |
| oligoGT375 | GAAGACAATGCCGGAGAGAGACCCAGTACCA GTAGGGCAGTGAGCGCAAC | Fw primer for constructing pGT270 |
| oligoGT376 | GAAGACTTTAGTAGTATGAGACCGGAAAGGA AACTATGCGGCATCAGAGC | Rv primer for constructing pGT270 |
| oligoGT436 | TATGTCAGTAAAAAAAAAGGAGGTTCGTTAT GTGAAGAGCGTAAGACCTCTAGGGCGGCG | Fw primer for cloning RBSc4 into pStA0 as a composite part |
| oligoGT437 | TACGAAAGATTTACGAAAGGAGGTTGGTTAT GTGAAGAGCGTAAGACCTCTAGGGCGGCG | Fw primer for cloning RBSc21 into pStA0 as a composite part |
| oligoGT438 | AGTGTGTTCCAAGAGAAAGGAGGTATCTGAT GTGAAGAGCGTAAGACCTCTAGGGCGGCG | Fw primer for cloning RBSc48 into pStA0 as a composite part |
| oligoGT439 | TAGGTATATGCAATCAAAGGAGGTATATGAT GTGAAGAGCGTAAGACCTCTAGGGCGGCG | Fw primer for cloning RBSc110 into pStA0 as a composite part |
| oligoGT440 | TGTGTAGGTCCCACCAAAGGAGGTCTGTAAT GTGAAGAGCGTAAGACCTCTAGGGCGGCG | Fw primer for cloning RBSc123 into pStA0 as a composite part |
| oligoGT441 | GGCGCGAGTCAACGTTCCTGAGGAGTGGCAT GTGAAGAGCGTAAGACCTCTAGGGCGGCG | Fw primer for cloning RBSc249 into pStA0 as a composite part |
| oligoGT442 | AGGTATATTATACTGTTGGGGCAGTTACTTG TCAAGGTCATCTGGCTGTGAAGAGCCACACT GGATTCTCACCAATAAAAAACGC | Rv primer for cloning SPLc3 into pStA0 as a composite part |
| oligoGT443 | AACTAAATTATACTGATATTTAGTTAAATTG TCAATACGAGCCGCCTGTGAAGAGCCACACT GGATTCTCACCAATAAAAAACGC | Rv primer for cloning SPLc19 into pStA0 as a composite part |
| oligoGT444 | ACCTTAATTATATCAGAGCGCAGAGTGCTTG TCAATTTTTCGTGGCTGTGAAGAGCCACACT GGATTCTCACCAATAAAAAACGC | Rv primer for cloning SPLc25 into pStA0 as a composite part |

| | | |
|---|---|---|
| oligoGT445 | ACGTTTATTATATAGTTAGCGGGCCGCTGTGTCAAAGGTGGCGAGCTGTGAAGAGCCACACTGGATTCTCACCAATAAAAAACGC | Rv primer for cloning SPLc45 into pStA0 as a composite part |
| oligoGT446 | TATTCCATTATATCTCGGTAACTCCTGGCTGTCAATAGACGTCACCTGTGAAGAGCCACACTGGATTCTCACCAATAAAAAACGC | Rv primer for cloning SPLc47 into pStA0 as a composite part |
| oligoGT578 | CGGCGGGTTTTTTTATAGCTAAAAGGACGAAGAGCGACCTCTAGGGCGGCGG | Fw primer for cloning ECK120015170 into pStA0 |
| oligoGT579 | AAGCGGGTTTTTTCGAAAATTGTTTATGAAGAGCCACACTGGATTCTCACCAATAAAAAACGC | Rv primer for cloning ECK120015170 into pStA0 |
| oligoGT583 | /5Phos/CGAATCATTATAGGGTTGTGTTCTCTGCGTGTCAAAGCTTAGGGGCTGTGAAGAGCCACACTGGATTCTCACCAATAAAAAACGC | Rv primer for cloning SPLc17 into pStA0 as a composite part |
| oligoGT603 | CCATAAAGCCAACCACACCT | Rv primer to screen for presence of lycopene pathway in *ssl0410* locus |
| oligoGT605 | TTGATCGAACGGGGTGTTAT | Fw primer to screen for presence of lycopene pathway in *ssl0410* locus |
| oligoGT653 | GATCTGGCATGGGTGGAG | Rv primer to screen for presence of lycopene pathway in *ssl0410* locus |
| oligoGT654 | ACCCTGAATGATGCTCTCCA | Fw primer to screen for presence of lycopene pathway in *ssl0410* locus |
| oligoAH34 | CAGGAAATATTGCTTTGCAGG | Fw sequencing primer for pATM2 and all derivative plasmids |
| oligoAH35 | ACACCCCTTGTATTACTGTTTATGT | Rv sequencing primer for pATM2 and all derivative plasmids |
| oligoAH48 | ACTTTGGTTGCTACCTCGCT | Fw primer to screen for integration into *ssl0410* locus |
| oligoAH49 | AGGGAAAATGCCAAAACGCA | Rv primer to screen for integration into *ssl0410* locus |

**Table S4. Table of plasmids used in this study.**

| Plasmid | Relevant properties or Description | Source |
|---|---|---|
| pATM2 | pSHUTTLE2 derivative with promoter Psll1120 and EYFP reporter | [2] |
| pATM10 | pSHUTTLE2 derivative with promoter PpsbA2 and EYFP reporter | This study |
| pStA0 | Start-Stop Assembly empty storage vector | [1] |
| pStA1AB | Start-Stop Assembly Level 1 vector (A and B fusion sites) | [1] |
| pStA1BC | Start-Stop Assembly Level 1 vector (B and C fusion sites) | [1] |
| pStA1CD | Start-Stop Assembly Level 1 vector (C and D fusion sites) | [1] |
| pStA1DZ | Start-Stop Assembly Level 1 vector (D and Z fusion sites) | [1] |
| pStA1DZ | Start-Stop Assembly Level 1 vector (D and Z fusion sites) | [1] |
| pStA212 | Start-Stop Assembly Level 2 vector (1 and 2 fusion sites) | [1] |
| pGT270 | pSHUTTLE2 derivative with pStA212 Start-Stop Assembly cassette | This study |
| pGT287 | pStA0::SPLc3-RBSc4 (composite promoter-RBS part) | This study |
| pGT288 | pStA0::SPLc19-RBSc4 (composite promoter-RBS part) | This study |
| pGT289 | pStA0::SPLc25-RBSc4 (composite promoter-RBS part) | This study |
| pGT290 | pStA0::SPLc45-RBSc4 (composite promoter-RBS part) | This study |
| pGT293 | pStA0::SPLc3-RBSc21 (composite promoter-RBS part) | This study |
| pGT294 | pStA0::SPLc19-RBS21 (composite promoter-RBS part) | This study |
| pGT295 | pStA0::SPLc25-RBSc21 (composite promoter-RBS part) | This study |
| pGT296 | pStA0::SPLc45-RBSc21 (composite promoter-RBS part) | This study |
| pGT297 | pStA0::SPLc47-RBS21 (composite promoter-RBS part) | This study |
| pGT300 | pStA0::SPLc19-RBS48 (composite promoter-RBS part) | This study |
| pGT301 | pStA0::SPLc25-RBSc48 (composite promoter-RBS part) | This study |
| pGT302 | pStA0::SPLc45-RBSc48 (composite promoter-RBS part) | This study |
| pGT303 | pStA0::SPLc47-RBSc48 (composite promoter-RBS part) | This study |
| pGT305 | pStA0::SPLc3-RBSc110 (composite promoter-RBS part) | This study |
| pGT306 | pStA0::SPLc19-RBSc110 (composite promoter-RBS part) | This study |
| pGT307 | pStA0::SPLc25-RBSc110 (composite promoter-RBS part) | This study |

| | | |
|---|---|---|
| pGT308 | pStA0::SPLc45-RBSc110 (composite promoter-RBS part) | This study |
| pGT309 | pStA0::SPLc47-RBSc110 (composite promoter-RBS part) | This study |
| pGT311 | pStA0::SPLc3-RBSc123 (composite promoter-RBS part) | This study |
| pGT312 | pStA0::SPLc19-RBSc123 (composite promoter-RBS part) | This study |
| pGT313 | pStA0::SPLc25-RBSc123 (composite promoter-RBS part) | This study |
| pGT314 | pStA0::SPLc45-RBSc123 (composite promoter-RBS part) | This study |
| pGT315 | pStA0::SPLc47-RBSc123 (composite promoter-RBS part) | This study |
| pGT317 | pStA0::SPLc3-RBSc249 (composite promoter-RBS part) | This study |
| pGT318 | pStA0::SPLc19-RBSc249 (composite promoter-RBS part) | This study |
| pGT319 | pStA0::SPLc25-RBsc249 (composite promoter-RBS part) | This study |
| pGT320 | pStA0::SPLc45-RBsc249 (composite promoter-RBS part) | This study |
| pGT321 | pStA0::SPLc47-RBSc249 (composite promoter-RBS part) | This study |
| pGT338 | pStA0::L3S2P21 | [1] |
| pGT339 | pStA0:: ECK120033737 | [1] |
| pGT340 | pStA0:: ECK120029600 | [1] |
| pGT356 | pStA0::*crtI* | [1] |
| pGT357 | pStA0::*crtB* | [1] |
| pGT358 | pStA0::*crtE* | [1] |
| pGT359 | pStA0::*dxs* | [1] |
| pGT424 | pStA0::ECK120015170 | This study |
| pGT425 | pStA0::SPLc17-RBSc4 (composite promoter-RBS part) | This study |
| pGT426 | pStA0::SPLc17-RBSc21 (composite promoter-RBS part) | This study |
| pGT427 | pStA0::SPLc17-RBSc48 (composite promoter-RBS part) | This study |
| pGT428 | pStA0::SPLc17-RBSc110 (composite promoter-RBS part) | This study |
| pGT429 | pStA0::SPLc17-RBSc123 (composite promoter-RBS part) | This study |
| pGT430 | pStA0::SPLc17-RBSc249 (composite promoter-RBS part) | This study |
| pGT432 | High expression design for lycopene pathway | This study |
| pGT433 | Medium expression design for lycopene pathway | This study |
| pGT434 | Low burden design for lycopene pathway | This study |

| | | |
|---|---|---|
| pGT435 | High burden design for lycopene pathway | This study |
| pGT437 | Lycopene pathway clone randomly chosen from pGT547 | This study |
| pGT438 | Lycopene pathway clone randomly chosen from pGT547 | This study |
| pGT439 | Lycopene pathway clone randomly chosen from pGT547 | This study |
| pGT440 | Lycopene pathway clone randomly chosen from pGT547 | This study |
| pGT441 | Lycopene pathway clone randomly chosen from pGT547 | This study |
| pGT442 | Lycopene pathway clone randomly chosen from pGT547 | This study |
| pGT443 | Lycopene pathway clone randomly chosen from pGT547 | This study |
| pGT444 | Lycopene pathway clone randomly chosen from pGT547 | This study |
| pGT445 | Lycopene pathway clone randomly chosen from pGT547 | This study |
| pGT446 | Lycopene pathway clone randomly chosen from pGT547 | This study |
| pGT447 | Lycopene pathway clone randomly chosen from pGT547 | This study |
| pGT448 | Lycopene pathway clone randomly chosen from pGT547 | This study |
| pGT449 | Lycopene pathway clone randomly chosen from pGT547 | This study |
| pGT450 | Lycopene pathway clone randomly chosen from pGT547 | This study |
| pGT451 | Lycopene pathway clone randomly chosen from pGT547 | This study |
| pGT452 | Lycopene pathway clone randomly chosen from pGT547 | This study |
| pGT453 | Lycopene pathway clone randomly chosen from pGT547 | This study |
| pGT454 | Lycopene pathway clone randomly chosen from pGT547 | This study |
| pGT455 | Lycopene pathway clone randomly chosen from pGT547 | This study |
| pGT456 | Lycopene pathway clone randomly chosen from pGT547 | This study |
| pGT457 | Lycopene pathway clone from pGT547 with prescreen | This study |
| pGT458 | Lycopene pathway clone from pGT547 with prescreen | This study |
| pGT459 | Lycopene pathway clone from pGT547 with prescreen | This study |
| pGT460 | Lycopene pathway clone from pGT547 with prescreen | This study |
| pGT461 | Lycopene pathway clone from pGT547 with prescreen | This study |
| pGT462 | Lycopene pathway clone from pGT547 with prescreen | This study |
| pGT463 | Lycopene pathway clone from pGT547 with prescreen | This study |
| pGT464 | Lycopene pathway clone from pGT547 with prescreen | This study |

| | | |
|---|---|---|
| pGT465 | Lycopene pathway clone from pGT547 with prescreen | This study |
| pGT466 | Lycopene pathway clone from pGT547 with prescreen | This study |
| pGT467 | Lycopene pathway clone from pGT547 with prescreen | This study |
| pGT468 | Lycopene pathway clone from pGT547 with prescreen | This study |
| pGT469 | Lycopene pathway clone from pGT547 with prescreen | This study |
| pGT470 | Lycopene pathway clone from pGT547 with prescreen | This study |
| pGT471 | Lycopene pathway clone from pGT547 with prescreen | This study |
| pGT472 | Lycopene pathway clone from pGT547 with prescreen | This study |
| pGT473 | Lycopene pathway clone from pGT547 with prescreen | This study |
| pGT474 | Lycopene pathway clone from pGT547 with prescreen | This study |
| pGT475 | Lycopene pathway clone from pGT547 with prescreen | This study |
| pGT476 | Lycopene pathway clone from pGT547 with prescreen | This study |
| pGT517 | Lycopene pathway clone from pGT547 with prescreen | This study |
| pGT518 | Lycopene pathway clone from pGT547 with prescreen | This study |
| pGT519 | Lycopene pathway clone from pGT547 with prescreen | This study |
| pGT520 | Lycopene pathway clone from pGT547 with prescreen | This study |
| pGT547 | Lycopene pathway library | This study |

**Table S5. List of synthetic DNA sequences used in the study**

| Synthetic DNA sequence name | Sequence (5'-3') |
| --- | --- |
| *crtI* recoded (1647 bp) | atgacatcagctctcccccgcccggcaccaagtccgtacgcacgccgtaaaac<br>ggcgttggttattggcgcaggttttggtgggttggccctgggcattcgtctac<br>agtcgttaggttttgatacaacaattttggaacgtctggatggtcctggtggt<br>cgcgcgtatcaaaaacgtaccccagatggctatgtctttgacatgggtccgac<br>tgtgctgacggtgccgcattttatcgaagaactgtttgcgcttgaacgtgatc<br>gtgccggcctggatgcccccgattatcctcctgaagtgttgtcgggcgagcgc<br>gttaaggaaggcgtttctggtggcccgcatacgagccggtatgtcaccttagt<br>gccgattctgccctttaccgcattgtttttcacgatggcacgtattttgatt<br>atgatggcgaccctgaaagtactcggcggcagattgctgaattggcccctggc<br>gacttagccgggtatgaacgctttcatgccgatgccgaggccatctttcgtcg<br>gggcttcctggaactgggctacacgcactttggtgacgtcccgacgatgctgc<br>gggttgtacctgatctgctcaaactggacgccgttcggaccctgttctccttt<br>acgagtaagtactttcagagcgacaaactgcgccaagtgttctcttttgaaac<br>ccttttggtgggtgggaatcctctgagtgtgccggcgatctatgcaatgattc<br>acttcgttgaaaagacttgggggatccactatgccatgggcggcacaggcgca<br>ctggtgcgcggcctagtccaaaaatttgaggagctgggtggcgccattcgtta<br>tggcgccggcgtcgatgaagtactggtggatggcaatctgcctggtaaacgca<br>cagcgcggggtgtgcgcctggaaagcggcgaagaactgcgcgccgacctggtg<br>gcgtccaatggcgattgggctaacacgtatctgaaacgcgtccggccatcggc<br>acgtctggtcaactctgatttacgcgtgaaagccgcatctgaaagtatgagcc<br>tgctcgtggtttatttcgggtttcgcggcggtgatgacctgcccctcaaacat<br>cataatattttattaggcccacgctacgaggctctgctgagcgaaatctttgg<br>cacaaaacggttgggcgaagattttagccagtacctgcacgtcccaacgctca<br>ccgatccggctctggcacccgcgggtcatcatgcggcctatacacttgtcccg<br>gtgccgcataatggctcgggtattgattgggacgtggaaggtccaaagcttgc<br>cgaagcagccctggcagatatcgagcgccgcggtttgattccgggcctccgtg<br>aacggctcacacattttgaatttattacgccagattatttcgcaggcactctc<br>gattcctatctggggaacgcgtttggtccggagccgcgtctggtccagtcggc<br>atttttccgcccgcacaaccgcagcgaggatctccacaactttacttagtcg<br>gggcgggcgcgcagccaggcgcaggcacaccgagcgttatgatgtccgcgaaa<br>atgacagcgcgcctaatcgctgaagatttcggtatccatgctgatatccggcg<br>ctaa |

*crtB*
recoded
(978 bp)

Atgcgtagtcgcgctggtctgagcttacggttacccacgcgtaccttgaccgt
gaccgattactcccccgccctgccctgcaccgaactgcgccgtcctccactgg
ctcaggcggttcgctactgtcgggatttgacccgccagcactcaaagaccttc
tatctgggttcacagctcttttcgcctccggaacgcgcggcagtttgggcagt
gtatgcggcgtgccgcgcgggcgatgacatcgtcgatgaagccggcaacggcg
accgcgaacgcgaattgcgggaatggcgcagccgtattgatgccgcgtttgct
ggccaaccagcggatgatcccatctcaaccgcgctggcctgggcggcaggtcg
gtacgccatcccgcactcagctttcgcggaactgcatgaaggcctcaacatgg
atttacgcggtcatgaataccgtgatatggatgacttgttactgtattgccgc
cgtgtggcaggtgtggttggctttatggtggcaccgatttctggctaccgtgg
gggggctgctaccctgaatgatgctctccaactagggcaggcgatgcaactga
cgaatattctgcgcgatgtcggtgaagatctgacccgcggccgcgtatacctg
ccacagtctctgcttgatgaatatggcctgtctcgcgccgcgttagagcgctg
gggtcagggtgagcccctgtcaccggcctaccgtgctctcatgactcatcttg
gcggccttgcacgtgaatggtatgcagcaggtcgtgctggtattcctcaactt
gatggacgcggtcctctcgccgttctgactgccgcccgtgcgtatgagggtat
tctggacgatttggaacgggccggctacgacaacttcggtcggcgcgcgtacg
tgtcaggtcgtcgtaaacttctgatgttaccgcaggcctggtgggaactgcgt
agtctgggcgctgtccacggctaa

*crtE*
recoded
(990 bp)

atgcgcccggaattactcgcacgcgtgttaagcctgttaccggaaacctccgc
gacgccggaattggcacgcttttacgcgctcctgcgcgactatcctcaacgtg
gtggcaagggcattcggtcagaattactgcttgcctctgctcgtgcgcacggc
ctgtccgagtcagataccggttgggagtcagcattatggctggcggcagcctt
agaactgtttcagaactgggtgctggtgcacgatgatattgaagatgattcgg
aagaacgccgtggtcgtccggccctgcaccacttgtgtggtatgccggtcgct
cttaacgtgggggacgcgctgcacgcttacatgtgggctgctgttgggaaagc
caatgttccgggagcgtttgaagagtttctgcagatggtgtaccgcacggcgg
aaggccagcatctggatctggcatgggtggagggtcgtgaatggggcctgcgt
cccgcggattatctccagatggttggcctgaaaaccgcacactacacggttat
cgtgccgttacgtctgggggccctggcggcaggcatggcaccgcaggacgcgt
tcaccccagcgggtctggcgctgggtaccgcgtttcagattcgtgacgatgtc
ctcaatctggcaggtgatccggtgaagtatggtaaagaaattggtggcgatct
gttggaaggtaaacgtactctgattgtcctggactggttgactacggcgccgg
atgatcgcaaagccatcttcctggaccagatgcgtcggcaccgcgcagataaa
gaccctgcggtgatcgatgaaattcaccgctggctgcttgaaagcggctctgt
ggaagcggcgcaggactacgcgcaggcacaagccgcggaaggtctggacttgc
ttgaaaaagcattggcagacgcgccggatgcccaggccgccgctgccttactc
gcttctgttcgggaactggccacccgcgaaaaataa

*dxs*
recoded
(1863 bp)

atgtcttttgatattgcgaaatatccaaccctggccctagttgactcgactca
ggaattacgcctgctgccgaaagagagccttccaaagctgtgcgatgagttac
gccgctacctcctggattctgttagtcgtagctccggccacttcgcatcggga
ctaggtaccgtcgaactgacggttgcactgcattatgtgtataacaccccgtt
cgatcaattaatttgggacgtcggtcatcaagcatatccgcataaaattctga

ccggtcgtcgcgacaagatcggcacaattcgtcaaaaaggtggattgcatcct
ttcccgtggcgcggcgagtccgaatacgatgtgttgagcgttggccattcgtc
aacttctatcagcgctggtattggtattgcggtcgccgcggagaaagaaggca
aaaatcggcgtacggtttgcgtgatcggggatggtgcaatcaccgcaggcatg
gcattcgaagcgatgaaccatgcgggggacattcgtccggatatgctagtgat
tctgaacgataacgaaatgagtatttccgagaacgtgggggctcttaataacc
acttagcgcagctgctgagcggtaaactttattctagcctgcgcgaaggcggt
aagaaagtgttctcaggcgtccctccgattaaagagctccttaagcgtactga
ggaacacattaagggtatggtggttccaggcaccctgttcgaagaactgggtt
ttaattacattggtcctgtggacggccatgacgtgttaggcttaattaccacg
ttaaaaaacatgcgcgatctgaaaggacctcagtttctccacatcatgaccaa
gaaaggtcgcggctatgaaccggccgagaaagatccgattaccttccacgcag
ttccgaaattcgacccatcctccggctgtctgccgaaaagcagcggaggcctg
ccgagctattccaaaatctttggtgactggctgtgcgaaactgcggcaaaaga
taacaaactgatggccatcaccccggcaatgcgcgaaggttcgggaatggtgg
aattctctcgcaagttcccagaccgttattttgacgttgctatcgccgagcaa
catgcagtaacctttgcggctggtctggcaattggtggttataagccaattgt
ggccatctattcaacgttcttacaacgtgcatatgaccaggtgctacatgatg
tggccattcagaaactgccggtcttatttgccatcgatcgtgcggggatcgtt
ggggcagatggtcagacgcatcagggcgccttcgacctaagttatctgcgctg
tatcccagaaatggtgattatgacccctagcgatgagaatgaatgtcgtcaga
tgttatacaccggctatcattacaacgacggaccatccgctgtgcgctacccg
cgcggaaacgcggtgggcgttgaactgaccccgttagaaaaattaccgatcgg
aaaagggattgttaaacgtcgcggagaaaaactcgctattctgaacttcggca
cacttatgcccgaagcggctaaagtagcggaaagcctaaacgcaaccttggtg
gatatgcgcttcgtaaagccactggacgaagccctaatcttagagatggccgc
gtcacatgaggcgttagttacggtagaggaaaacgctattatgggcggagcgg
gtagtggtgtgaacgaagttctgatggctcaccgcaaaccggtgccggtactg
aacatcggcctcccggatttttcattccccaaggtactcaagaggaaatgcg
cgccgagctcggcttagatgctgcaggaatggaagccaaaattaaggcctggc
ttgcataa

**Table S6. Promoters and RBSs stored in pStA0.** Fw primer and Rv primer signify the oligonucleotides used for the inverse PCR carried out to generate the given construct.

| Plasmid ID | Promoter | RBS | Fw primer | Rv primer |
|---|---|---|---|---|
| pGT287 | SPLc3 | RBSc4 | oligoGT436 | oligoGT442 |
| pGT288 | SPLc19 | RBSc4 | oligoGT436 | oligoGT443 |
| pGT289 | SPLc25 | RBSc4 | oligoGT436 | oligoGT444 |
| pGT290 | SPLc45 | RBSc4 | oligoGT436 | oligoGT445 |
| pGT293 | SPLc3 | RBSc21 | oligoGT437 | oligoGT442 |
| pGT294 | SPLc19 | RBSc21 | oligoGT437 | oligoGT443 |
| pGT295 | SPLc25 | RBSc21 | oligoGT437 | oligoGT444 |
| pGT296 | SPLc45 | RBSc21 | oligoGT437 | oligoGT445 |
| pGT297 | SPLc47 | RBSc21 | oligoGT437 | oligoGT446 |
| pGT300 | SPLc19 | RBSc48 | oligoGT438 | oligoGT443 |
| pGT301 | SPLc25 | RBSc48 | oligoGT438 | oligoGT444 |
| pGT302 | SPLc45 | RBSc48 | oligoGT438 | oligoGT445 |
| pGT303 | SPLc47 | RBSc48 | oligoGT438 | oligoGT446 |
| pGT305 | SPLc3 | RBSc110 | oligoGT439 | oligoGT442 |
| pGT306 | SPLc19 | RBSc110 | oligoGT439 | oligoGT443 |
| pGT307 | SPLc25 | RBSc110 | oligoGT439 | oligoGT444 |
| pGT308 | SPLc45 | RBSc110 | oligoGT439 | oligoGT445 |
| pGT309 | SPLc47 | RBSc110 | oligoGT439 | oligoGT446 |
| pGT311 | SPLc3 | RBSc123 | oligoGT440 | oligoGT442 |
| pGT312 | SPLc19 | RBSc123 | oligoGT440 | oligoGT443 |
| pGT313 | SPLc25 | RBSc123 | oligoGT440 | oligoGT444 |
| pGT314 | SPLc45 | RBSc123 | oligoGT440 | oligoGT445 |
| pGT315 | SPLc47 | RBSc123 | oligoGT440 | oligoGT446 |
| pGT317 | SPLc3 | RBSc249 | oligoGT441 | oligoGT442 |
| pGT318 | SPLc19 | RBSc249 | oligoGT441 | oligoGT443 |
| pGT319 | SPLc25 | RBSc249 | oligoGT441 | oligoGT444 |
| pGT320 | SPLc45 | RBSc249 | oligoGT441 | oligoGT445 |

| | | | | |
|---|---|---|---|---|
| pGT321 | SPLc47 | RBSc249 | oligoGT441 | oligoGT446 |
| pGT425 | SPLc17 | RBSc4 | oligoGT436 | oligoGT583 |
| pGT426 | SPLc17 | RBSc21 | oligoGT437 | oligoGT583 |
| pGT427 | SPLc17 | RBSc48 | oligoGT438 | oligoGT583 |
| pGT428 | SPLc17 | RBSc110 | oligoGT439 | oligoGT583 |
| pGT429 | SPLc17 | RBSc123 | oligoGT440 | oligoGT583 |
| pGT430 | SPLc17 | RBSc249 | oligoGT441 | oligoGT583 |
